# PyINETA: Open-source platform for INADEQUATE-JRES integration in NMR metabolomics

**DOI:** 10.1101/2024.07.10.601875

**Authors:** Rahil Taujale, Mario Uchimiya, Chaevien S. Clendinen, Ricardo M. Borges, Christoph W. Turck, Arthur S. Edison

## Abstract

Annotating compounds with high confidence is a critical element in metabolomics. ^13^C-detection NMR experiment INADEQUATE (incredible natural abundance double-quantum transfer experiment) stands out as a powerful tool for structural elucidation, whereas this valuable experiment is not often included in metabolomics studies. This is partly due to the lack of community platform that provides structural information based INADEQUATE. Also, it is often the case that a single study uses various NMR experiments synergistically to improve the quality of information or balance total NMR experiment time, but there is no public platform that can integrate the outputs of INADEQUATE and other NMR experiments either. Here, we introduce PyINETA, Python-based INADEQUATE network analysis. PyINETA is an open-source platform that provides structural information of molecules using INADEQUATE, conducts database search, and integrates information of INADEQUATE and a complementary NMR experiment ^13^C *J*-resolved experiment (^13^C-JRES). Those steps are carried out automatically, and PyINETA keeps track of all the pipeline parameters and outputs, ensuring the transparency of annotation in metabolomics. Our evaluation of PyINETA using a model mouse study showed that our pipeline successfully integrated INADEQUATE and ^13^C-JRES. The results showed that ^13^C-labeled amino acids that were fed to mice were transferred to different tissues, and, also, they were transformed to other metabolites. The distribution of those compounds was tissue-specific, showing enrichment of particular metabolites in liver, spleen, pancreas, muscle, or lung. The value of PyINETA was not limited to those known compounds; PyINETA also provided fragment information for unknown compounds. PyINETA is available on NMRbox.

---

Robust annotation of compounds is a critical task in metabolomics. In NMR metabolomics, compound annotation is primarily based on chemical shifts of ^1^H, ^13^C, or both. Two-dimensional (2D) experiments increase the confidence level of annotation, providing correlations between protons or protons and carbons in molecules^1^. Although ^1^H-detection 2D experiments have been successfully implemented in metabolomics for this purpose, ^2, 3^ ^13^C-detection NMR can complement ^1^H-NMR and improve the quality of information^4, 5^. ^13^C-NMR has a broader chemical shift range and fewer overlapping peaks than ^1^H-NMR, which is ideal for metabolomics samples that are complex mixtures of compounds^6^. ^13^C-NMR can directly detect quaternary carbons, which leads to a broader coverage of carbon information in molecules. Most importantly from a perspective of structural elucidation, ^13^C-NMR can directly extract the backbone structure of molecules^7, 8^, essential information in structural elucidation.

Among various ^13^C-NMR experiments, INADEQUATE (incredible natural abundance double-quantum transfer experiment)^9^ stands out as a powerful tool for structural elucidation. This experiment unambiguously detects ^13^C-^13^C connectivity and extracts networks of carbons in molecules. INADEQUTE suffers from low natural abundance of ^13^C-^13^C couplings in molecules (*i.e.*, less than 1 in every 10^4^ C-C bonds), but this experiment can benefit from isotopic enrichment and becomes applicable to metabolomics samples^8^. Although one could apply INADEQUATE to many samples in a metabolomics study and profile the metabolome differences between samples^8^, INADEQUATE requires a relatively long time for data collection, and this approach is not always practical especially when spectrometer time is limited. Thus, it is useful to have a profiling experiment that requires less instrument time but can be easily used with INADEQUATE. Although an obvious choice is a simple 1D ^13^C experiment to profile all samples in a study, ^13^C-enriched samples leads to complicated peak shapes and more overlap than experiments at natural abundance ^13^C.

Our approach to the problem is to use a 2D ^13^C *J*-resolved experiment (^13^C-JRES), which separates chemical shifts from coupling constants into different dimensions. A 1D projection of a 2D ^13^C-JRES is free from multiplets and can be collected quickly enough for efficient profiling. The output can be statistically processed^10^ and linked to 2D spectra for a representative sample such as internal pooled sample, a mixture of aliquots of all the study samples, for annotation. This can meet the requirement of less overall experiment time and reliable compound information. It has been shown that a combination of ^13^C-JRES for profiling and INADEQUATE for annotation can achieve both reducing overall experiment time and maintaining the quality of structural information for a metabolomics study^11^.

In addition to the robust compound annotation, the benefit of introducing INADEQUATE is also its suitability for computational tasks. Clendinen *et al*. (2015) developed INETA (INADEQUATE network analysis) that computationally constructs networks of backbone carbons in molecules using the INADEQUATE rules. In INADEQUATE, two directly bonded carbon atoms resonate at their natural frequencies along the acquisition dimensions and at the sum of their frequencies along the indirect double-quantum dimension. This leads to pairs of peaks that are symmetric along a diagonal with slope 2 (*i.e.*, *Y* = 2*X* on an INADEQUATE spectrum), and these pairs of INADEQUATE peaks are then linked vertically to expand the network. INETA used the constructed networks to search an internal INADEQUATE database, which was simulated using assigned ^13^C chemical shifts and chemical structures of compounds deposited in Biological Magnetic Resonance Bank (BMRB)^12^. INETA was used to annotate the endo- and exometabolomes of ^13^C-enriched *Caenorhabditis elegans*^8^.

Despite the clear advantages of being able to annotate metabolites using INADEQUATE, there are several obstacles to its routine use. First, samples need to be isotopically labeled with ^13^C. Many microorganisms and plants can be uniformly enriched using a carbon source such as ^13^C-glucose at modest cost^5^. This is more challenging for human studies, but select targeted pathways using isotope tracers with *ex vivo* tissue slices or cell cultures are regularly studied^13–15^. This paper applied the approach in isotopically-labeled mice to show feasibility. Second, access to a high-sensitivity cryogenic ^13^C NMR probe is necessary. Such probes are made commercially and can be accessed through large NMR facilities with user programs such as The National High Magnetic Field Lab (https://nationalmaglab.org/) or The Network for Advanced NMR (https://usnan.nmrhub.org/). Third, the previous software developed by our group to perform INETA was written using Mathematica, which is not open source. Finally, software has not previously been developed to integrate INADEQUATE with ^13^C JRES data.

Here, we propose PyINETA, an open-source platform that can automatically integrate INADEQUATE and JRES data. In addition to the functions that were originally implemented in INETA, our new PyINETA seamlessly transfers INADEQUATE information to JRES, providing compound information to individual JRES peaks. The pipeline is run on Python, and researchers can freely implement this open-source platform to various metabolomics studies. We evaluated the applicability of PyINETA using a model mouse study, in which metabolites originating from ^13^C-labeled diet were examined.

As the number of metabolomics publications increases rapidly, transparency of studies is becoming more critical than ever^16^. This is especially true for compound annotation where the basis for annotation is required with significant rigor^17^ but is not always reported in publication^18^. PyINETA is designed to report all the annotation steps, providing a community platform that ensures the reproducibility and transparency of compound annotation in metabolomics.

## Experimental Section

We first developed PyINETA, a Python package that is capable of annotating compounds based on INADEQUATE and transferring the compound information to ^13^C-JRES. We then demonstrate the functionality by evaluating a variety of tissues collected from a mouse study where mice were fed with a diet that contained ^13^C-labled amino acids.

### Development of PyINETA

PyINETA performs a series of tasks, including importing data, constructing networks, database matching for INADEQUATE spectra, and transferring the annotation information to JRES peaks. The pipeline requires two input file types, configuration file and spectra. Configuration file contains all the information relating to parameters used for the analysis. Available PyINETA parameters are summarized in Supplementary Table S1. After a configuration file and INADEQUATE and JRES spectra are loaded, the pipeline initializes a PyINETA class object. This object contains two chemical shift vectors for a spectrum (*i.e.*, ppm values for the direct and double-quantum dimensions), along with an intensity matrix. The input spectra are .ft files prepared by NMRPipe^19^.

Next, PyINETA defines peaks. First, the pipeline collects peak data points from a spectrum. For this step, users define the maximum and minimum thresholds for intensity (parameters ‘PPmax’ and ‘PPmin’, respectively), and those values are used to distinguish signals from background noise. To collect data points, the pipeline initially uses a threshold of PPmax. Then, it iteratively decreases the threshold and collect data points until the threshold reaches PPmin. Users can define the number of iterations (‘steps’). Any data points collected on a previous iteration step are not subjected to subsequent steps. To define peaks from collected data points, clustering was used. For each iteration step, collected data points are first clustered along the direct dimension using a threshold for the distance of the center of mass (‘PPCS’). Then, data points are clustered along the double-quantum dimension using a second threshold (‘PPDQ’). Each cluster represents a single peak. Their center of mass corresponds to a peak center, and all other points define peak area.

Once peaks are defined, the pipeline creates INADEQUATE networks. In INADEQUATE, carbons next to each other in a molecule have the same chemical shift value in the double-quantum dimension. Using this rule, horizontally aligned peaks are extracted. To meet the definition of horizontally aligned peaks, the difference in chemical shift between peaks in the double-quantum dimension needs to be less than a threshold (parameter ‘DQT’). INADEQUARTE has another rule that, for carbons next to each other in a molecule, the sum of chemical shift values in the direct dimension is equal to their chemical shift values in the double-quantum dimension (*i.e.*, sum-rule). Using this rule, horizontally aligned peaks are screened:

|(CS1 + CS2) − mean [DQ1,DQ2]| <= SumXY

where CS1 and CS2 represent chemical shift values for Carbons 1 and 2 in the direct dimension, DQ1 and DQ2 chemical shift values in the double-quantum dimension for Carbons 1 and 2, SumXY a threshold, respectively. Also, in INADEQUTE, carbons next to each other in a molecule have the same distance from a line of *Y* = 2*X* (*i.e.*, diagonal-rule). Horizontal peaks are further screened by the diagonal-rule to define horizontal networks:

|(CS1-DQ1)/2 - (CS2-DQ2)/2|<= SDT

where SDT is a threshold. When multiple horizontal networks are originating from sequential carbons in a single compound, those horizontal networks can be linked by vertically aligned peaks of a shared carbon. To meet the definition of vertically aligned peaks, two peaks need to have a chemical shift difference less than a threshold (‘CST’) in the direct dimension.

Once networks are created, PyINETA starts database matching. In this pipeline, every single peak in the network is initially compared to peaks in a simulation database in the PyINETA package. This simulated database is created using experimental spectra deposited in BMRB^12, 20^ (details are in Section ‘*Generation of a simulated INADEQUATE database*’). Firstly, when the distance between sample peaks and database peaks is less than a threshold (‘CSMT’), peaks are considered as matched peaks. Secondly, when the number of chemical shift matches between database networks and sample networks is more than a threshold (‘NCMT’), they are considered to be matched networks. Matched networks are subsequently analyzed along the double-quantum dimension. For this step, when the difference in chemical shift between database networks and sample networks is less than a threshold (‘DQMT’), database networks are considered as matched networks. Since all the database entries that satisfy the criteria are considered as matched networks in this scheme, it is potentially possible to find multiple matches for any given network. To evaluate the resulting matches, two scores, hit score and coverage score, are assigned to each matched network. Hit score quantifies the proportion of peaks in sample networks that matched a specific database entry, whereas coverage score represents the proportion of peaks in a database entry that matched those in a sample network. For both hit score and coverage score, 1 is the maximum value. For JRES peaks, peak area values are calculated (‘Peak_Width_1D’) and the presence and absence of peaks corresponding to INADEQUATE are defined using a threshold value (‘Intensity_threshold_1D’).

Finally, a summary file reports all the major statistics about the number of peaks that passed every step. Results from each step are also saved as pickle files (*i.e.*, an object serialization mechanism in Python).

The developed pipeline is installed on NMRbox^21^. The source code is also available on GitHub (https://github.com/edisonomics/PyINETA.git), along with instructions and example datasets.

### Generation of a simulated INADEQUATE database

We constructed a simulated INADEQUATE database using structural information and experimental 1D ^13^C spectra deposited in the BMRB database^12^. The simulated database we used in this study contains 1,973 entries, covering 1,209 metabolites. The simulated database is available as a json file in the package. Additionally, when users need to create their own simulated spectra, the module gen_PyINETAdb.py is available. Input format for this module is either NMR-STAR^22^ or tables with chemical shifts and structural information. The ambiguity information for peaks in input files are retained as ambiguity scores and reported in the final PyINETA output.

### In vivo ^13^C labeling and sample preparation

All animal experiments and protocols have been reviewed and approved by the Institutional Animal Care and Use Committee of the Max Planck Institute of Psychiatry. Three 8-week-old male C57BL/6 mice (Charles River Laboratories, Maastricht, The Netherlands) were housed under standard conditions (12-h light/dark cycle, lights on at 0600 h, room temperature 23 ± 2 °C, humidity 60%, tap water and food *ad libitum*) and fed with standard rodent diet (Harlan Laboratories, Inc., Indianapolis, IN, USA) for one week. For adaptation prior to labeling the animals were first fed an unlabeled *Ralstonia eutropha* bacterial protein-based rodent diet (Silantes GmbH, Munich, Germany) for 4 days. The food supply was then switched to ^13^C-labeled *Ralstonia eutropha* bacterial diet (Silantes GmbH) for 14 days (Supplementary Tables S2 and S3). Following labeling the animals were sacrificed and organs and blood isolated. The partially ^13^C-labeled animals did not show any discernible health effects compared to animals fed with a standard diet, and had similar weight gains as animals fed with standard food (data not shown). Tissues were homogenized in 30 volumes of ice cold 80% methanol, homogenates were centrifuged and supernatants were dried. Dried samples were resuspended in 50 µL of deuterated water and methanol (1:4 volume ratio) (Supplementary Table 4).

### NMR data collection and processing

Data were collected on a Bruker Avance Neo 900 MHz with a 5-mm TXO cryoprobe (Bruker), using NMR tubes with a diameter of 1.7 mm (Bruker). For INADEQUATE experiment, default pulse programs with adiabatic 180° pulses (Bruker nomenclature, inadphppsp) was used. For JRES, Bruker’s default pulse program (jresdcqf) was modified to implement an adiabatic 180° pulse after we verified adiabatic 180° pulse is necessary in collecting ^13^C-JRES on our 900 MHz magnet^11^. Detailed parameter settings are in Supplementary Table S5. TopSpin 4.0.9 was to operate the spectrometer.

All the NMR spectra were processed using NMRPipe^19^. Briefly, for both INADEQUATE and JRES, FID was Fourier-transformed after applying a squared sine-bell function and a double zero-filling on both direct and indirect dimensions. For JRES, spectra were further tilted and symmetrized. Detailed NMRPipe processing parameters are in Supplementary Table S6. Further data processing for JRES spectra was conducted using Metabolomics Toolbox (https://github.com/edisonomics/metabolomics_toolbox) on MATLAB R2022b (MathWorks). Briefly, projection spectra were created from JRES, and they were aligned with the CCOW method (function ‘guide_align1D’) and normalized by the probabilistic quotient normalization (PQN) method^23^ (‘normalize’). The ALATIS numbering system^24^ was used for describing carbon numbers. All the raw data, NMRPipe processing scripts, processed data, PyINETA output files, and MATLAB scripts are available in Metabolomics Workbench with Study ID ST003304.

## Results and Discussion

### PyINETA provides a flexible environment for INADEQUATE-JRES integration

Since PyINETA uses various parameters to perform a series of tasks, we utilize a system of ‘configuration file’ (Figure 1, left), which contains all the parameters that will be used in the pipeline (Supplementary Information 1 for an example configuration file). Users can manage and overview the pipeline with this single stage.

**Figure 1.**
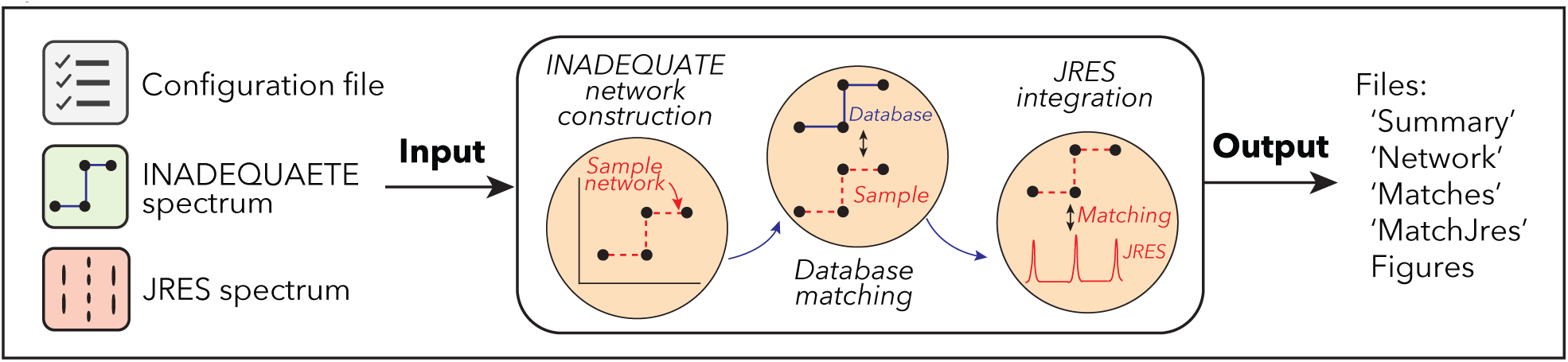
Workflow of PyINETA

Users can also fine tune each step using this configuration file. For example, signal-to-noise levels in JRES can vary greatly between samples, and users can set threshold parameters that are appropriate for a specific spectrum. In optimizing parameters, users do not need to run the whole pipeline, and each step can be run separately by defining the -s option. This saves computational time.

Using parameters in a configuration file and an input INADEQUATE spectrum (Supplementary Figure S1-a), the pipeline constructs INADEQUATE networks and searches for the constructed networks in an internal database (Figure 1, middle). The internal database is based on ^13^C chemical shift and chemical structure of metabolites deposited in BMRB^12^, which was originally implemented in Clendinen et al. (2015). BMRB is one of the largest databases in experimental NMR data from small molecule metabolomics. When users have a compound(s) of interest that are not deposited in BMRB, they can manually add those compounds to the internal PyINETA database. This includes other experimental or computational databases that provide ^13^C peaks assigned to a known structure and computed chemical shifts from putative compounds.

### PyINETA Workflow

Finally, as a key component, PyINETA integrates the information of INADEQUATE and ^13^C-JRES (Figure 1, middle). PyINETA reads a raw ^13^C-JRES spectrum and creates a projection. Then, peaks are picked from the projection spectrum, peaks between INADEQUATE and JRES are matched, and compound information based on INADEQUATE will be transferred to JRES. We made this component optional so that users can still use this pipeline when only INADEQUATE spectra are available.

After this processing, results are reported as a set of output files (Figure 1, right); ‘Summary file’ overviews the number of peaks passed every step in INADEQUATE processing steps (Supplementary Information 2 for an example file). ‘Networks file’ provides chemical shift values for network peaks (Supplementary Information 3). ‘Matches file’ shows matched database entries, compound names, and peak connectivity information (Supplementary Information 4). Matches file also contains confidence scores for those matches (*i.e.*, ambiguity score, hit score, and coverage score; see Experimetal Section for details), and users can evaluate the reliability of annotation. The INADEQUATE-JRES integration step is summarized as an output file ‘file_5MatchJres.xlsx’. (Supplementary Information S5).

In addition to those summary files, PyINETA provides figures for individual networks and matched compounds for INADEQUATE (examples will follow in the next section). Similarly, PyINETA creates figures for matched peaks for INADEQUATE and JRES. This capability was implemented to enable users to further validate the results of the automatic annotation. The annotation information made by this new PyINETA is consistent with that of the original study INETA^8^ (Supplementary Information 6; Supplementary Table S7).

### 13C-JRES profiling showed clear tissue-specific spectral patterns

We applied ^13^C-JRES profiling to a mouse study where a fate of a diet was investigated (Figure 2a). Three mice were fed with a diet that contained ^13^C-labeled amino acids, including nine essential and seven essential amino acids. After this feeding, mouse tissues were collected and the distribution of metabolites originating from those amino acids were analyzed in several tissues including liver, adrenal gland, lung, muscle, pancreas, plasma, brain, spleen, and thymus. JRES spectra in those tissues were consistent between the three mice (Supplementary Figure S2), and the higher intensities were observed in liver samples. Also, ^13^C-JRES peaks were composed of sharp singlets, as expected.

**Figure 2.**
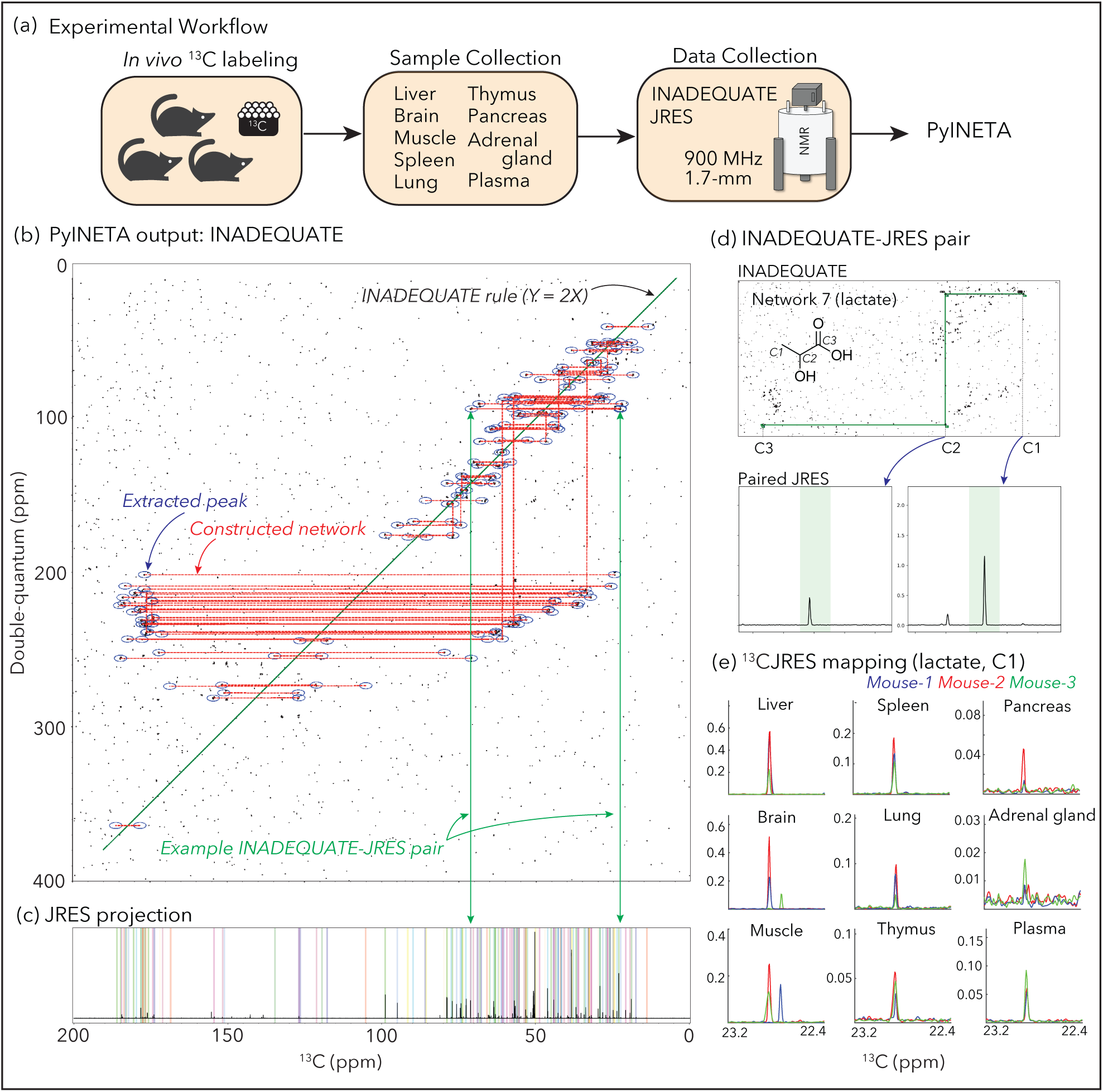
(a) Experimental workflow of this study; (b) Example of INADEQUATE networks for a mouse liver sample constructed by PyINETA; (c) JRES projection for the same sample. Peaks picked by PyINETA are highlighted in different colors; (d) INADEQUATE-JRES pair found by PyINETA. A compound name was also assigned using a database, which is lactate for this pair; (e) Mapping of a JRES peak in different tissues. JRES peaks for C1 for lactate are shown here. All the plots for (b), (c), and (d) are from the original outputs from PyINETA, with a slight graphical modification. Chemical structure was drawn using ChemDraw V23.

### PyINETA was able to integrate INADEQUATE and JRES information automatically

Since the profiling results was consistent between mice, and the liver samples had the highest intensities, we first used one of the liver samples (Sample ID, 30) for the evaluation of INADEQUATE-JRES integration in PyINETA. From an INADEQUATE spectrum collected for the liver sample, PyINETA constructed 67 INADEQUATE networks (Figure 2b; Supplementary Information 3 for a list of all networks). The majority (52 out of 67) of the networks were single network that linked two peaks, whereas 15 networks were longer, containing 8 peaks at maximum in a network (Network 8) (Supplementary Information 3 for a complete list). Networks with just two carbons could reflect the original structure of compound, including the case where networks are fragmented spectroscopically by heteroatoms in molecules. They, however, could also originate from compounds with longer-backbones when whole networks were not created computationally and fragmented into shorter networks. We observed both cases in our output. For example, choline is a compound with a backbone of a single network (C-C) and was found in a single horizontal network of Network 44 (Supplementary Figure S1-b). On the other hand, lactate, which has a network of three carbons (C-C-C), was found in two separate horizontal networks (Networks 7 and 55) because of missing vertical connection of two horizontal networks (Supplementary Figures S1-c and d). Those broken networks can be manually inspected or improved by relaxing the tolerance parameter for vertical network construction (CST; Supplementary Table S1).

PyINETA then searched for those 67 networks in the database, and 46 networks matched with at least one candidate compound (Supplementary Information 4). The rest of the 21 networks did not have any matched compound, indicating that they are compounds that are not in BMRB (‘unknown compound’ hereafter).

One network can potentially match more than one compound in the database when compound structures are similar. Also, a single compound can potentially exist in more than one network as described above. We further investigated the results using PyINETA’s function of output figures and excluded matches with less confidence due to partial structural similarity. As a result, the matched networks were those for 21 compounds (Table 1). They included amino acids (alanine, glutamine, leucine, lysine, threonine, glutamic acid, isoleucine, valine, and proline), an amino sugar (D_glucuronate), an amino alcohol (2_Aminoethyl_dihydrogen_phosphate), amino sulfonic acids (hypotaurine and taurine), a pyrimidine (barbituric acid), amines (betaine, choline, ethanolamine, and putrescine), and organic acids (lactic acid and chloroacetic acid). They also included 16 unknown compounds (Table 1).

**Table 1.**
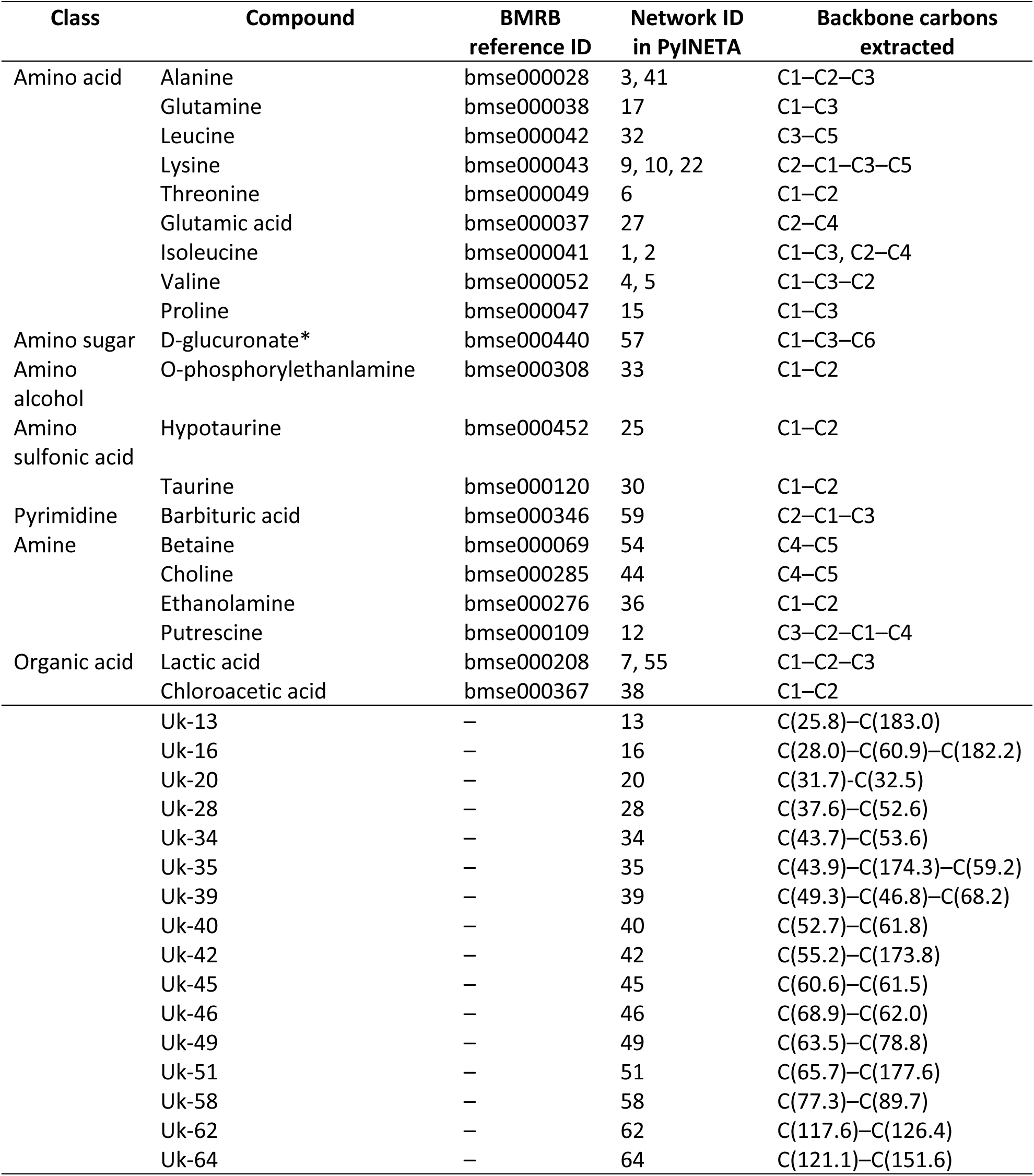
List of compounds and corresponding INADEQUATE networks detected by PyINETA for a mouse liver sample. Only a conservative list of metabolites based on the inspection of the original output is shown. The Original outputs are in Supplementary Information 3 and 4. For numbering of carbons, the ALATIS numbering system was used, except for unknown compounds where chemical shift values were indicated in parentheses. *Only partial structural information was available in the BMRB entry.

Next, PyINETA analyzed a JRES spectrum collected for the same sample (Supplementary Figure S1-e). Among 67 INADEQUATE networks, 65 of them had JRES peaks in the corresponding regions (Figure 2c; Supplementary Information S5 for a complete list). PyINETA then transferred compound information based on INADEQUATE to JRES peaks (Figure 2d). PyINETA seamlessly paired INADEQUATE and JRES.

### PyINETA revealed the distribution of metabolites originating from a diet in different tissues

Since PyINETA transformed compound information to JRES peaks, we were able to examine the distribution of a specific metabolite in different tissues based on JRES (Figure 2e). We further extended this analysis to other metabolites and examined the distribution of metabolites in different tissues. For this analysis, we used representative JRES peaks (no overlapping peaks with a minimum intensity of 0.1), and 19 compounds are included in the following analysis. We found three different categories (Figure 3). Among the 19 compounds, 13 of them were enriched in liver compared with other tissues (Compound Type-A) (Figure 3). Compounds in this category are amino acids (lysine, glutamic acid, alanine, and glutamine), an organic acid (lactic acid), and an amino sugar (D-glucuronate). On the other hand, two metabolites were depleted in liver but enriched in other tissue(s) (Type-B) (Figure 3); they included an amino alcohol (O-phosphorylethanlamine) in pancreas and spleen, and an amino sulfonic acid (hypotaurine) in pancreas. Finally, four compounds are enriched in both liver and other tissues (muscle, spleen, lung, or pancreas) (Type-C; Figure 3). They are amino acids (valine, threonine, and isoleucine) and an amino sulfonic acid (taurine). Since INADEQUATE detects ^13^C-^13^C coupling in molecules that occurs less than 0.01% in natural abundance, here we interpret that the metabolites in our results are originating from the ^13^C that were fed to the mice, and the effects of natural abundance metabolites that were originally present in tissues are negligible.

**Figure 3.**
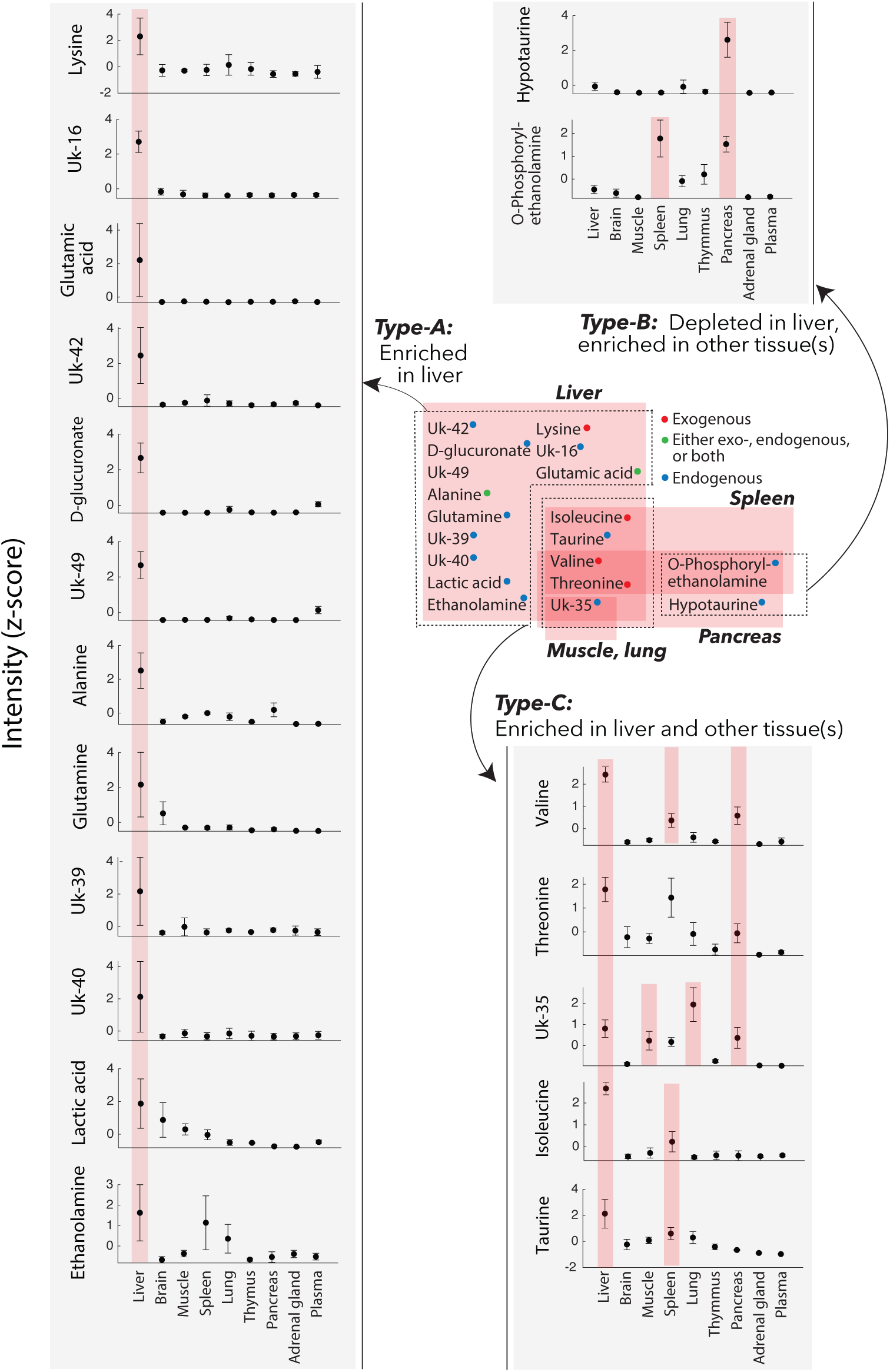
Distribution of compounds in the mouse tissues. Those are compounds originated from ^13^C-labeled diet the mice were fed with. Gray insets: Peak intensities based on JRES (z-scored). Error bars, standard deviation (*n* = 3). When a compound in a specific tissue is enriched compared with any other tissue, they are highlighted in pink (ANOVA with multiple comparison; a complete statistical summary is in Supplementary Figure S3. The values for this figure are also in Metabolomics Workbench).

There could be different sources explaining those metabolites. Lysine, isoleucine, valine, and threonine are essential amino acid and were included in the ^13^C-labeled amino acids in the diet, suggesting that those amino acids were directly distributed from the diet to tissues (*i.e.*, exogenous; Supplementary Figure S4). Alanine and glutamic acid were also contained in the diet, but, also, they are non-essential amino acids and the mice might have synthesized them (*i.e.*, endogenous), leaving a possibility that those amino acids were either exogenous, endogenous, or both (Supplementary Figure S4). On the other hand, glutamine, another non-essential amino acid was not included in the diet, indicating that glutamine was exclusively endogenous in this system. Metabolites other than proteinogenic amino acids (D-glucuronate, lactic acid, ethanolamine, taurine, O-phosphorylethanol amine, hypotaurine), they are exclusively exogenous (Supplementary Figure S4).

Liver plays a central role in amino acid metabolism, and this was reflected by our results. Net uptake of alanine predominantly occurs in liver^25^, consistent with the observation of enriched alanine in liver (Type-A pattern). Alanine is further used in liver to produce other metabolites including glutamate^26^, which was also in the Type-A pattern. Alanine also serves as a major precursor for gluconeogenesis which occurs in liver^26, 27^. Those transformation processes suggest that the rate of alanine uptake was exceeding that of transformation, resulting in the enriched alanine observed in this study. Similarly, liver is one of the dominant tissues that take up glutamine^26, 27^. On the other hand, branched-chain amino acids (BCAAs) isoleucine, valine, and threonine were not exclusively high in liver but were also abundant in other tissues (Type-B). This could be due to the fact that BCAAs can escape catabolism in liver because of low activity of BCAA transferases and inefficient uptake of BCAAs in liver^25, 27, 28^. Other compounds abundant in liver are also related to those metabolism and roles. Lactate is a precursor for gluconeogenesis which occurs in liver^29^, taurine one of the abundant amino acids with diverse physiological functions in liver^30^, and glucuronic acid a compound that is used in glucuronidation in liver^31^. On the contrary, hypotaurine was depleted in liver but enriched in pancreas. High level of hypotaurine biosynthesis occurs in pancreas in mouse^32^.

### PyINETA also revealed the distribution of unknown compounds originating from the diet

PyINETA was useful even when compounds are not in the database. Out of 67 networks, 21 did not match any of entry in BMRB. Even under that situation, we were able to track those compounds, showing its backbone structure and distribution in different tissues (Figure 3; Table 1). For example, Uk-16 is a compound that is not in the database, but PyINETA extracted its backbone structure and chemical shift information. Because INADEQUATE and JRES are already linked by PyINETA, we were able to trace this unknown compound using JRES and revealed the distribution in different tissues. Uk-16 was exclusively enriched in liver compared with other tissues, indicating that this is a compound that is actively processed in liver but has not been covered in the BMRB.

In NMR metabolomics in general, database matching is primarily focused on peaks that matched database compounds, and peaks that did not match database compounds are usually not retained. On the other hand, PyINETA treats matched and unmatched networks equally and provides structural information. Since PyINETA has a capability of adding new entries to the internal database, users can make use of the obtained knowledge on unknown compounds in future studies. If there is a match of an unknown compound between studies, that is a finding of a common unknown compound.

Tracking metabolites using stable isotopes to understand metabolic pathway and flux has been an active field since its establishment^13, 33^. Despite the success and tremendous value of this approach to track targeted compounds^14^, investing unknown compounds in this framework is a laborious task, and effort has been made to develop untargeted approaches regardless of analytical platform^34^. PyINETA is capable of handling unknown compounds and can contribute to tackling this challenge in this field.

### Ensuring the reproducibility of compound annotation in metabolomics

PyINETA keeps track of all the parameters used in the pipeline as a configuration file. Also, the results from individual steps are saved as pickle files. Those pickle files contain all the information of the PyINETA class that is required to reproduce the results. Because of this system, annotation information from PyINETA is completely reproducible. Users can also deposit those files to databases such as Metabolomics Workbench^35^ along with original data to ensure the reproducibility of compound annotation in a study.

## Conclusion

PyINETA removes current stumbling blocks in the field of metabolomics, making the best use of ^13^C-NMR and improving the transparency and reproducibility of compound annotation. In addition to the example study we presented here, PyINETA is expandable to any system that can be labeled to address specific questions in various research fields. PyINETA is installed on NMRbox^21^ and publicly available.

## Supporting information

Taujale_et_al_supplementary_Information

## Supporting Information

Additional experimental details, tables, and figures (S1-S4) are provided separately as a .docx file.

## Acknowledgements

We thank John Glushka for providing suggestions on INADEQUATE and JRES experiments and Laura Morris and NMRbox staff members for installing PyINETA on NMRbox. Christopher S. Esselman provided valuable feedback on PyINETA. This project was supported by NSF Network for Advanced NMR (NAN) (1946970) and an NIH MIRA award 5R35GM148240 awarded to A.S.E. and the Max Planck Society (C.W.T.).

## Author contribution

R.T. and M.U. contributed equally to this work. A.S.E. and C.T. designed the study; R.T. developed PyINETA; C.W.T. supervised mouse experiments and sample collection; C.S.C. prepared the mouse samples for NMR; M.U. conducted the NMR experiments; M.U., R.T., R.M.B., and A.S.E. analyzed the data. M.U., R.T., and A.S.E wrote the manuscript with all authors’ input.

## Data availability

PyINETA is available on NMRbox. The source code, example datasets, and instructions are also available on GitHub (https://github.com/edisonomics/PyINETA.git). All the raw NMR data, NMRPipe processing scripts, processed data, PyINETA output files, and MATLAB scripts used in this study are deposited to Metabolomics Workbench with Study ID ST003304.

## References

1. Bingol, K.; Bruschweiler, R., Multidimensional approaches to NMR-based metabolomics. Anal Chem 2014, 86 (1), 47–57.

2. Bingol, K.; Li, D. W.; Zhang, B.; Bruschweiler, R., Comprehensive metabolite identification strategy using multiple two-dimensional NMR spectra of a complex mixture implemented in the COLMARm web server. Anal Chem 2016, 88 (24), 12411–12418.

3. Bhinderwala, F.; Vu, T.; Smith, T. G.; Kosacki, J.; Marshall, D. D.; Xu, Y.; Morton, M.; Powers, R., Leveraging the HMBC to Facilitate Metabolite Identification. Anal Chem 2022, 94 (47), 16308–16318.

4. Edison, A. S.; Le Guennec, A.; Delaglio, F.; Kupce, E., Practical guidelines for ^13^C-based NMR metabolomics. Methods Mol Biol 2019, 2037, 69–95.

5. Clendinen, C. S.; Stupp, G. S.; Ajredini, R.; Lee-McMullen, B.; Beecher, C.; Edison, A. S., An overview of methods using ^13^C for improved compound identification in metabolomics and natural products. Front Plant Sci 2015, 6, 611.

6. Clendinen, C. S.; Lee-McMullen, B.; Williams, C. M.; Stupp, G. S.; Vandenborne, K.; Hahn, D. A.; Walter, G. A.; Edison, A. S., ^13^C NMR metabolomics: Applications at natural abundance. Anal Chem 2014, 86 (18), 9242–50.

7. Bingol, K.; Zhang, F.; Bruschweiler-Li, L.; Bruschweiler, R., Carbon backbone topology of the metabolome of a cell. J Am Chem Soc 2012, 134 (21), 9006–11.

8. Clendinen, C. S.; Pasquel, C.; Ajredini, R.; Edison, A. S., ^13^C NMR metabolomics: INADEQUATE network Analysis. Anal Chem 2015, 87 (11), 5698–706.

9. Bax, A.; Freeman, R.; Kempsell, S. P., Natural abundance ^13^C-^13^C coupling observed via double-quantum coherence. J Am Chem Soc 1980, 102 (14), 4849–4851.

10. Robinette, S. L.; Lindon, J. C.; Nicholson, J. K., Statistical spectroscopic tools for biomarker discovery and systems medicine. Anal Chem 2013, 85 (11), 5297–303.

11. Uchimiya, M.; Olofsson, M.; Powers, M. A.; Hopkinson, B. M.; Moran, M. A.; Edison, A. S., ^13^C NMR metabolomics: J-resolved STOCSY meets INADEQUATE. J Magn Reson 2023, 347, 107365.

12. Ulrich, E. L.; Akutsu, H.; Doreleijers, J. F.; Harano, Y.; Ioannidis, Y. E.; Lin, J.; Livny, M.; Mading, S.; Maziuk, D.; Miller, Z.; Nakatani, E.; Schulte, C. F.; Tolmie, D. E.; Wenger, R. K.; Yao, H. Y.; Markley, J. L., BioMagResBank. Nuc Acids Res 2008, 36, D402–D408.

13. Bartman, C. R.; Faubert, B.; Rabinowitz, J. D.; Deberardinis, R. J., Metabolic pathway analysis using stable isotopes in patients with cancer. Nat Rev Cancer 2023, 23 (12), 863–878.

14. Lin, P. H.; Lane, A. N.; Fan, T. W. M., Stable isotope-resolved metabolomics by NMR. Nmr-Based Metabolomics: Methods and Protocols 2019, 2037, 151–168.

15. Hattori, A.; Tsunoda, M.; Konuma, T.; Kobayashi, M.; Nagy, T.; Glushka, J.; Tayyari, F.; Cskimming, D. M.; Kannan, N.; Tojo, A.; Edison, A. S.; Ito, T., Cancer progression by reprogrammed BCAA metabolism in myeloid leukaemia. Nature 2017, 545 (7655), 500-+.

16. Wilkinson, M. D.; Dumontier, M.; Aalbersberg, I. J.; Appleton, G.; Axton, M.; Baak, A.; Blomberg, N.; Boiten, J. W.; da Silva Santos, L. B.; Bourne, P. E.; Bouwman, J.; Brookes, A. J.; Clark, T.; Crosas, M.; Dillo, I.; Dumon, O.; Edmunds, S.; Evelo, C. T.; Finkers, R.; Gonzalez-Beltran, A.; Gray, A. J.; Groth, P.; Goble, C.; Grethe, J. S.; Heringa, J.; t Hoen, P. A.; Hooft, R.; Kuhn, T.; Kok, R.; Kok, J.; Lusher, S. J.; Martone, M. E.; Mons, A.; Packer, A. L.; Persson, B.; Rocca-Serra, P.; Roos, M.; van Schaik, R.; Sansone, S. A.; Schultes, E.; Sengstag, T.; Slater, T.; Strawn, G.; Swertz, M. A.; Thompson, M.; van der Lei, J.; van Mulligen, E.; Velterop, J.; Waagmeester, A.; Wittenburg, P.; Wolstencroft, K.; Zhao, J.; Mons, B., The FAIR Guiding Principles for scientific data management and stewardship. Sci Data 2016, 3, 160018.

17. Sumner, L. W.; Amberg, A.; Barrett, D.; Beale, M. H.; Beger, R.; Daykin, C. A.; Fan, T. W.; Fiehn, O.; Goodacre, R.; Griffin, J. L.; Hankemeier, T.; Hardy, N.; Harnly, J.; Higashi, R.; Kopka, J.; Lane, A. N.; Lindon, J. C.; Marriott, P.; Nicholls, A. W.; Reily, M. D.; Thaden, J. J.; Viant, M. R., Proposed minimum reporting standards for chemical analysis Chemical Analysis Working Group (CAWG) Metabolomics Standards Initiative (MSI). Metabolomics 2007, 3 (3), 211–221.

18. Powers, R.; Andersson, E. R.; Bayless, A. L.; Brua, R. B.; Chang, M. C.; Cheng, L. L.; Clendinen, C. S.; Cochran, D.; Copié, V.; Cort, J. R.; Crook, A. A.; Eghbalnia, H. R.; Giacalone, A.; Gouveia, G. J.; Hoch, J. C.; Jeppesen, M. J.; Maroli, A. S.; Merritt, M. E.; Pathmasiri, W.; Roth, H. E.; Rushin, A.; Sakallioglu, I. T.; Sarma, S.; Schock, T. B.; Sumner, L. W.; Takis, P.; Uchimiya, M.; Wishart, D. S., Best practices in NMR metabolomics: current state. TrAC Trends in Analytical Chemistry 2024, 171.

19. Delaglio, F.; Grzesiek, S.; Vuister, G. W.; Zhu, G.; Pfeifer, J.; Bax, A., NMRPipe - A multidimensional spectral processing system based on Unix pipes. J Biomol Nmr 1995, 6 (3), 277–293.

20. Hoch, J. C.; Baskaran, K.; Burr, H.; Chin, J.; Eghbalnia, H. R.; Fujiwara, T.; Gryk, M. R.; Iwata, T.; Kojima, C.; Kurisu, G.; Maziuk, D.; Miyanoiri, Y.; Wedell, J. R.; Wilburn, C.; Yao, H. Y.; Yokochi, M., Biological Magnetic Resonance Data Bank. Nucleic Acids Res 2023, 51 (D1), D368–D376.

21. Maciejewski, M. W.; Schuyler, A. D.; Gryk, M. R.; Moraru, I. I.; Romero, P. R.; Ulrich, E. L.; Eghbalnia, H. R.; Livny, M.; Delaglio, F.; Hoch, J. C., NMRbox: A resource for biomolecular NMR computation. Biophys J 2017, 112 (8), 1529–1534.

22. Ulrich, E. L.; Baskaran, K.; Dashti, H.; Ioannidis, Y. E.; Livny, M.; Romero, P. R.; Maziuk, D.; Wedell, J. R.; Yao, H.; Eghbalnia, H. R.; Hoch, J. C.; Markley, J. L., NMR-STAR: comprehensive ontology for representing, archiving and exchanging data from nuclear magnetic resonance spectroscopic experiments. J Biomol Nmr 2019, 73 (1-2), 5–9.

23. Dieterle, F.; Ross, A.; Schlotterbeck, G.; Senn, H., Probabilistic quotient normalization as robust method to account for dilution of complex biological mixtures. Application in H-1 NMR metabonomics. Anal Chem 2006, 78 (13), 4281–4290.

24. Dashti, H.; Westler, W. M.; Markley, J. L.; Eghbalnia, H. R., Unique identifiers for small molecules enable rigorous labeling of their atoms. Sci Data 2017, 4, 170073.

25. Felig, P., Amino acid metabolism in man. Annu Rev Biochem 1975, 44, 933–955.

26. Bröer, S.; Bröer, A., Amino acid homeostasis and signalling in mammalian cells and organisms. Biochem J 2017, 474, 1935–1963.

27. Paulusma, C. C.; Lamers, W. H.; Broer, S.; van de Graaf, S. F. J., Amino acid metabolism, transport and signalling in the liver revisited. Biochem Pharmacol 2022, 201.

28. Bifari, F.; Nisoli, E., Branched-chain amino acids differently modulate catabolic and anabolic states in mammals: a pharmacological point of view. Brit J Pharmacol 2017, 174 (11), 1366–1377.

29. Gerich, J. E.; Meyer, C.; Woerle, H. J.; Stumvoll, M., Renal gluconeogenesis - Its importance in human glucose homeostasis. Diabetes Care 2001, 24 (2), 382–391.

30. Miyazaki, T.; Matsuzaki, Y., Taurine and liver diseases: a focus on the heterogeneous protective properties of taurine. Amino Acids 2014, 46 (1), 101–110.

31. Yang, G. Y.; Ge, S. F.; Singh, R.; Basu, S.; Shatzer, K.; Zen, M.; Liu, J.; Tu, Y. F.; Zhang, C. N.; Wei, J. B.; Shi, J.; Zhu, L. J.; Liu, Z. Q.; Wang, Y.; Gao, S.; Hu, M., Glucuronidation: Driving factors and their impact on glucuronide disposition. Drug Metab Rev 2017, 49 (2), 105–138.

32. Yoon, S. J.; Combs, J. A.; Falzone, A.; Prieto-Farigua, N.; Caldwell, S.; Ackerman, H. D.; Flores, E. R.; DeNicola, G. M., Comprehensive metabolic tracing reveals the origin and catabolism of cysteine in mammalian tissues and tumors. Cancer Res 2023, 83 (9), 1426–1442.

33. Bartman, C. R.; TeSlaa, T.; Rabinowitz, J. D., Quantitative flux analysis in mammals. Nat Metab 2021, 3 (7), 896–908.

34. Buckley, D. H.; Huangyutitham, V.; Hsu, S. F.; Nelson, T. A., Stable isotope probing with 15N achieved by disentangling the effects of genome G+C content and isotope enrichment on DNA density. Applied and Environmental Microbiology 2007, 73 (10), 3189–3195.

35. Sud, M.; Fahy, E.; Cotter, D.; Azam, K.; Vadivelu, I.; Burant, C.; Edison, A.; Fiehn, O.; Higashi, R.; Nair, K. S.; Sumner, S.; Subramaniam, S., Metabolomics Workbench: An international repository for metabolomics data and metadata, metabolite standards, protocols, tutorials and training, and analysis tools. Nucleic Acids Res 2016, 44 (D1), D463–D470.

