## Supplementary material for "PyINETA: Open-source platform for INADEQUATE-JRES integration in NMR metabolomics": Taujale_et_al_supplementary_Information: Taujale_et_al_supplementary_Information.pdf

Supplementary Information 1: Example of configuration file in PyINETA

Supplementary Information 2: Example of Summary file in PyINETA

Supplementary Information 3: Example of Networks file in PyINETA

Supplementary Information 4: Example of Matches file in PyINETA

Supplementary Information 5: Example of MatcheJRES file in PyINET

Supplementary Information 6: Evaluation of consistency of annotation between the original INETA and new PyINETA

***Supplementary Tables***

Supplementary Table S1. List of adjustable parameters in PyINETA

Supplementary Table S2. Composition of the diet used in this study

Supplementary Table S3. Composition of amino acids in the diet

Supplementary Table S4. Sample list of this study

Supplementary Table S5. Parameters used for NMR experiments.

Supplementary Table S6. Parameters used for NMRPipe processing

Supplementary Table S7. List of metabolites found by PyINETA for endo- and exometabolites from *Caenorhabditis elegans*

***Supplementary Figures***

Supplementary Figure S1: INADEQUATE and JRES spectra and PyINETA output figures from the mouse study

Supplementary Figure S2: JRES projection spectra for the mouse tissues

Supplementary Figure S3: Results of ANOVA and multiple comparison

Supplementary Figure S4: Possible sources of metabolites detected in this study

Supplementary Information 1: Example of configuration file in PyINETA.

```
; This is the template for a config.ini file used with pyINETA
; Lines enclosed in "[]" indicate section headers.
; Other lines have the options and parameters required by INETA
; Lines starting with a ";" are comments and describe the options
preceding it.

[PeakPick]
Ft_File =
<path_to_the_raw_data_file>/raw_data/pyINETA_AlaRef_2023/4_inad.ft2
Data_Matrix_File =
; (optional if no Ft file)
13C_Ppm_File =
; (optional if no Ft file)
Double_Quantum_File =
; (optional if no Ft file)

Xrange_min = 0
Xrange_max = 200
; The min and max ppm values for the X-axis (13C axis)
Yrange_min = 0
Yrange_max = 400
; The min and max ppm values for the Y-axis (DQ axis)
OutImage_pick_separate = fig_1plotSeparate.eps
OutImage_pick_complete = fig_1plotAll.eps
; Filenames for output images after peak picking

Shift = No
; Yes or No | Yes to shift the spectrum – This functionality is helpful
for aligning the INADEQUATE spectrum by shifting all the peaks uniformly
across one or both axes.
Direction = Pos
; Pos or Neg | Which direction to shift the spectra in
; For eg, Pos will move a peak at 35 ppm to 40 ppm
;         Neg will move a peak at 40 ppm to 35 ppm
Shift13C = 20
; # of units to shift the spectra
; In a typical INADEQUATE spectrum, 20 units ~ 1 ppm
Full13C = 4096
; Size of the 13C dimension in units; number of columns in the data
matrix.
FullDQ = 2048

; Size of the DQ dimension in units; number of rows in the data matrix.

PPmin = 5.7e10
; Minimum intensity to select a peak
PPmax = 4e11
; Maximum intensity of a peak
steps = 10
; Number of iterations of peak picking between PPmin and PPmax
; has to be at least 2
```

```
[ClusterPoints]
PPCS = 1
; All points within this range (in ppm) along the 13C axis will be
clustered into a single point.
;
PPDQ = 2
; All points within this range (in ppm) along the DQ axis will be
clustered into a single point.

OutImage_cluster_separate = fig_2clusterCenterSeparate.eps
OutImage_cluster_complete = fig_2clusterCenterAll.eps
; Filenames for output images after clustering

[FindNetwork]
Select = all
; all or last | use "all" for most cases
; "all" iterates over all steps to find peaks
; "last" selects peaks found in the last iteration only
LevelPointsDistance = 0.5
; Typical range: 0.5 - 2
; Distance between points (in ppm) across different levels to be
considered a new peak
; Lower values results in more points.
DQT = 0.2
; Typical range: 0.5 - 2
; (threshold for DQ - diffDQ < DQT for networking)
; (Control horizontal connection to be selected for network)
SumXY = 2
; Typical range: 0.5 - 1
; (threshold for sum 2X equals Y) (Control whether Y=X1+X2)
SDT = 0.5
; Typical range: 0.2 - 0.5
; (Symmetrical/Diagonal tolerance) (Equidistant from diagonal) 0.2-0.5
best if 0.3
CST = 0.05
; (chemical shift tolerance) (Controls vertical connections), 0.07

Network_output_file = file_3Networks.txt
OutImage_network_AllNets = fig_3findNetworkAllNets.eps

[MatchDatabase]
Database_file = <path_to_pyineta>/db/InetaDB.200922.json

Ambiguity = 1
; Higher values will include more database networks, for example, 0 – will
remove all, 1 – will include all
CSMT = 1
; 0.5,2 (chemical shift match tolerance,near) Higher value will increase
the x-axis range for a match between a network and db peak
Match_tolerance = 2
; NCMT, number of peak matches, Higher values will require more number of
matching peaks to consider a database hit
DQMT = 4
```

```
; (double quantum match tolerance) Higher value will increase the y-axis
range for a match between a network and db peak
Topology_tolerance = 2
; Higher values will require x and y values to be closer to the db peak to
consider a hit
Hit_Score_threshold = 0.2
; (ratio of matched-number of peaks in db) 0-1. A value of 1 means all db
peaks need to match.A value >1 indicates that multiple candidate peaks
matched the same db peak
Coverage_Score_threshold = 0.5
; (ratio of matched-number of peaks in network) 0-1. A value of 1 requires
all network peaks to match the db peaks

Matches_list_output_file = file_4Matches.txt
Summary_file = file_Summary.txt

[Overlay1D]
1D_File_List = <path_to_1D_file>/raw_data  /file1.ft,/file2.ft,/file3.ft
; Comma separated list of 1D ft filenames
Peak_Width_1D = 0.5
; Typical range: 0.2 - 2
; When matching INADEQUATE networks with 1D spectra, this value (in ppm)
determines the peak width used to calculate peak area.
; Range of peak width calculation will be (peakPostion-Peak_Width_1D/2 to
peakPostion+Peak_Width_1D/2)
Intensity_threshold_1D = 10000
; Typical range:
; Intensity threshold for area under the curve around matched peaks to
flag as present or absent.
Match1d_output_file = file_5Match1ds.txt
OutImage_Match1d = fig_5highlight1dMatches.png
; Filename for output file with 1D matching results.
OutImageFormat1D = "png"
; Output image format: can be eps, jpeg, jpg, pdf, pgf, png, ps, raw,
rgba, svg, svgz, tif, tiff

Shift_1D = No
; Yes or No | Yes to shift all the provided 1D spectra
Direction_1D = Neg
; Pos or Neg; Which direction to shift the spectra in
; For eg, Pos will move a peak at 35 ppm is now at 40 ppm
;         Neg will move a peak at 40 ppm is now at 35 ppm
Shift_1D_val = 930
; 930 - 1.42; 780 - 1.19 ppm
; # of units to shift the spectra (650 units ~ 1 ppm for a spectra fo size
131072 units)
Full_1D = 131072
; Size of the 13C dimension in units

[OverlayJres]
Jres_File_List = /<path_to_Jres_file>/3.tilt.sym.ft2
; 40_cres.tilt.sym.ft
; Comma separated list of Jres ft filenames.
```

```
Jres_Projection_Method = max
; Can be sum,max or avg
Peak_Width_Jres = 0.5
; Typical range: 0.2 - 2
; When matching INADEQUATE networks with 1D spectra, this value (in ppm) 
determines the peak width used to calculate peak area.
; Range of peak width calculation will be (peakPostion-Peak_Width_1D/2 to 
peakPostion+Peak_Width_1D/2)
Intensity_threshold_Jres = 10000
; Typical range: 
; Intensity threshold for area under the curve around matched peaks to 
flag as present or absent.
MatchJres_output_file = file_5MatchJres.txt
OutImage_MatchJres = fig_5highlightJresMatches.png
; Filename for output file with 1D matching results.
OutImageFormatJres = png
; Output image format: can be eps, jpeg, jpg, pdf, pgf, png, ps, raw, 
rgba, svg, svgz, tif, tiff

Shift_Jres = No
; Yes or No | Yes to shift all the provided 1D spectra
Direction_Jres = Neg
; Pos or Neg; Which direction to shift the spectra in
; For eg, Pos will move a peak at 35 ppm is now at 40 ppm
;         Neg will move a peak at 40 ppm is now at 35 ppm
Shift_Jres_val = 930
; 930 - 1.42; 780 - 1.19 ppm
; # of units to shift the spectra (650 units ~ 1 ppm for a spectra of size 
131072 units)
Full_Jres = 131072
; Size of the 13C dimension in units
```

Supplementary Information 2: Example of Summary file in PyINETA.

```
Summary for PyINETA run on 2024-06-07 15:17:21.483747 :
# of picked peaks in all steps: 6629
# of peaks after clustering: 2493
# of peaks retained after merging steps: 1806
# of networks found: 67
# of matches found: 46
```

Supplementary Information 3: Example of Networks file in PyINETA.

```
Network1    [27.03 40.91],[13.72 41.01]
Network2   [38.58 56.17],[17.27 55.93]
Network3   [18.7  72.18],[53.2  71.98]
Network4   [31.73 51.09],[19.2  50.97]
Network5   [31.82 52.36],[20.57 52.45]
Network6   [68.52 90.83],[22.24 90.78]
Network7   [22.87 94.29],[22.76 93.85],[70.98 94.12]
Network8   [24.7  51.54],[42.71 69.7 ],[ 42.75 107.44],[26.95
69.7 ],[ 64.76 107.56],[ 64.36 107.25],[26.95 51.62],[23.43 50.35],[26.95
50.46]
Network9   [24.13 53.52],[29.14 53.52]
Network10  [32.59 57.04],[24.3  56.98]
Network11  [ 24.64 201.46],[176.75 201.46]
Network12  [25.69 67.17],[41.51 67.17]
Network13  [183.03 208.87],[ 25.8  209.03]
Network14  [26.42 86.41],[60.42 86.62]
Network15  [48.6  75.22],[26.53 75.15]
Network16  [182.24 243.22],[28.07 89.06],[ 60.97 243.13],[60.99 89.21]
Network17  [28.95 85.85],[56.68 85.82]
Network18  [29.03 70.84],[ 33.61 213.39],[29.04 62.72],[179.7
213.2],[33.57 62.62],[41.66 71.  ]
Network19  [176.46 233.69],[29.51 86.95],[175.96 233.  ],[57.33
86.94],[ 57.33 233.61]
Network20  [32.53 64.2 ],[31.67 64.35]
Network21  [ 32.4  216.14],[183.72 216.19]
Network22  [32.76 89.8 ],[57.01 89.9 ]
Network23  [176.93 210.88],[ 34.1  210.98]
Network24  [39.26 75.41],[40.64 80.07],[39.28 80.07],[35.96 75.41]
Network25  [57.67 93.92],[36.09 94.03]
Network26  [183.33 219.95],[ 36.39 220.  ]
Network27  [ 36.72 221.34],[184.84 221.45]
Network28  [52.6  90.24],[37.61 90.11]
Network29  [ 37.96 214.07],[176.08 214.05],[176.12 238.9 ],[ 62.97 239.03]
Network30  [49.98 88.1 ],[38.15 88.13]
Network31  [41.77 97.25],[55.36 97.25]
Network32  [56.02 98.68],[42.53 98.68]
Network33  [ 43.35 106.54],[ 63.12 106.42]
Network34  [53.69 97.62],[43.76 97.62]
Network35  [ 44.   218.17],[ 43.95 218.75],[174.37 218.7 ],[174.43
218.33],[174.34 233.4 ],[ 59.26 233.4 ]
Network36  [ 60.23 104.39],[ 44.13 104.39]
Network37  [180.62 225.68],[ 44.86 225.79]
Network38  [178.15 223.78],[178.14 224.28],[ 46.   223.94],[ 46.14 224.34]
Network39  [ 46.86 115.12],[49.38 96.25],[46.84 96.37],[ 68.23 115.23]
Network40  [ 52.76 114.37],[ 61.89 114.52]
Network41  [ 53.12 230.86],[177.71 230.86]
Network42  [ 55.28 229.4 ],[173.87 229.56],[174.04 229.21]
Network43  [177.5  233.48],[ 55.83 233.72]
Network44  [ 58.28 128.55],[ 70.17 128.58]
Network45  [ 60.68 122.31],[ 61.52 122.37]
Network46  [ 68.97 130.51],[ 62.07 130.8 ],[ 61.04 130.4 ]
Network47  [175.75 238.53],[ 63.04 238.26]
```

```
Network48  [ 74.1  146.86],[ 72.59 146.83],[ 74.1  137.65],[ 63.49 137.64]
Network49  [ 63.56 142.6 ],[ 78.81 142.35]
Network50  [ 73.62 138.44],[ 64.94 138.43]
Network51  [ 65.74 243.31],[177.61 243.39]
Network52  [ 85.82 153.46],[ 67.59 153.56]
Network53  [ 73.8  142.03],[ 67.95 141.93]
Network54  [171.19 239.93],[ 68.8  239.96]
Network55  [ 71.04 255.69],[184.53 255.64]
Network56  [ 94.87 169.46],[ 74.43 150.18],[ 75.56 150.18],[ 74.41 169.4 ]
Network57  [ 78.72 155.7 ],[ 98.79 175.98],[ 76.99 155.7 ],[ 77.01 176.11]
Network58  [ 77.39 167.15],[ 89.73 167.02]
Network59  [ 79.69 251.91],[172.08 251.96]
Network60  [ 85.51 177.14],[ 91.48 177.14]
Network61  [168.57 273.8 ],[105.15 273.37]
Network62  [126.48 244.29],[117.68 244.4 ]
Network63  [119.39 254.13],[134.66 254.13]
Network64  [121.12 272.98],[151.69 272.98]
Network65  [126.82 281.2 ],[154.51 281.4 ]
Network66  [151.08 278.03],[126.96 278.13]
Network67  [178.26 363.89],[186.07 363.89]
```

Supplementary Information 4: Example of Matches file in PyINETA.

```
#NetworkNum ID MatchName  Solvent    AmbiguityScore Hitscore
CoverageScore  MatchedConnections UnmatchedConnections
Network1   bmse000041 L_isoleucine   D2O    0.0    0.2    1.0    CX1-CX2-
>C1-C6,
Network1   bmse000319 3_Methyl_2_oxopentanoic_acid   D2O    0.0    0.2
1.0    CX1-CX2->C5-C8,
Network1   bmse000574 3_Methyl_2_oxopentanoic_acid   D2O    0.0    0.2
1.0    CX1-CX2->C5-C8,
Network1   bmse000578 ethylmalonic_acid  D2O    0.0    0.25   1.0    CX1-
CX2->C6-C7,
Network1   bmse000866 L_isoleucine   D2O    0.0    0.2    1.0    CX1-CX2-
>C1-C6,
Network2   bmse000041 L_isoleucine   D2O    0.0    0.2    1.0    CX2-CX1-
>C7-C2,
Network2   bmse000866 L_isoleucine   D2O    0.0    0.2    1.0    CX2-CX1-
>C7-C2,
Network3   bmse000028 L_alanine  D2O    0.0    0.5    1.0    CX2-CX1->C2-
C5,
Network3   bmse000236 D_alanine  D2O    0.0    0.5    1.0    CX2-CX1->C2-
C3,
Network3   bmse000282 DL_Alanine D2O    0.0    0.5    1.0    CX2-CX1->C4-
C5,
Network3   bmse000403 Ala_Ala    D2O    0.0    0.25   1.0    CX2-CX1->C6-
C9,
Network4   bmse000052 L_valine   D2O    0.4    0.25   1.0    CX2-CX1->C7-
C5,
Network4   bmse000466 DL_beta_leucine    D2O    0.0    0.2    1.0    CX2-
CX1->C8-C4,
Network4   bmse000860 L_valine   D2O    0.4    0.25   1.0    CX2-CX1->C7-
C5,
Network5   bmse000052 L_valine   D2O    0.4    0.25   1.0    CX1-CX2->C5-
C7,
Network5   bmse000466 DL_beta_leucine    D2O    0.0    0.2    1.0    CX1-
CX2->C4-C8,
Network5   bmse000860 L_valine   D2O    0.4    0.25   1.0    CX1-CX2->C5-
C7,
Network6   bmse000049 L_threonine    D2O    0.0    0.333  1.0    CX2-CX1-
>C1-C2,
Network6   bmse000794 3_carboxypropyl_trimethyl_ammonium D2O    0.571
0.333  1.0    CX2-CX1->C5-C4,
Network6   bmse000859 L_threonine    D2O    0.0    0.333  1.0    CX2-CX1-
>C1-C2,
Network7   bmse000208 L_lactic_acid  D2O    0.0    0.5    0.667  CX3-CX1-
>C2-C3,CX3-CX3->C2-C2, CX2-CX3->?-?,
Network7   bmse000269 R_Lactate  D2O    0.0    0.5    0.667  CX3-CX1->C4-
C5,CX3-CX3->C4-C4, CX2-CX3->?-?,
Network7   bmse000979 L_lactic_acid  D2O    0.0    0.5    0.667  CX3-CX1-
>C2-C3,CX3-CX3->C2-C2, CX2-CX3->?-?,
Network9   bmse000043 L_lysine   D2O    0.0    0.2    1.0    CX1-CX2->C6-
C7,
Network9   bmse000072 cadaverine D2O    0.0    0.25   1.0    CX1-CX2->C7-
C6,
```

```
Network9   bmse000237 DL_pipecolic_acid  D2O    0.0    0.2    1.0    CX1-
CX2->C4-C5,
Network9   bmse000373 isovaleric_acid    D2O    0.0    0.25   1.0    CX1-
CX2->C6-C3,
Network9   bmse000411 L_Norleucine   D2O    0.0    0.2    1.0    CX1-CX2-
>C7-C5,
Network9   bmse000493 epsilon_caprolactone   CDCl3  0.0    0.2    1.0
CX1-CX2->C4-C3,
Network9   bmse000571 2_hydroxyhexanoic_acid D2O    0.0    0.2    1.0
CX1-CX2->C6-C4,
Network10  bmse000043 L_lysine   D2O    0.0    0.2    1.0    CX2-CX1->C6-
C5,
Network10  bmse000249 methyl_4_aminobutyrate D2O    0.0    0.333  1.0
CX2-CX1->C2-C4,
Network10  bmse000372 epsilon_caprolactam    D2O    0.0    0.2    1.0
CX2-CX1->C5-C3,
Network10  bmse000429 DL_2_Aminoadipic_acid  D2O    0.0    0.2    1.0
CX2-CX1->C7-C6,
Network10  bmse000442 L_homocitrulline   D2O    0.0    0.2    1.0    CX2-
CX1->C7-C9,
Network10  bmse000745 homoarginine   D2O    0.0    0.2    1.0    CX2-CX1-
>C7-C8,
Network11  bmse000145 N_acetyl_L_glutamine   D2O    0.286  0.2    1.0
CX1-CX2->C1-C2,
Network11  bmse000157 N_acetyl_L_alanine D2O    0.0    0.333  1.0    CX1-
CX2->C9-C7,
Network11  bmse000382 N_Acetyl_L_glutamic_acid   D2O    0.0    0.2    1.0
CX1-CX2->C13-C11,
Network11  bmse000423 N_acetyl_L_aspartic_acid   D2O    0.0    0.25   1.0
CX1-CX2->C12-C11,
Network11  bmse000468 4_Acetamidobutyric_acid    D2O    0.0    0.25   1.0
CX1-CX2->C10-C9,
Network11  bmse000707 N_acetyl_histidine D2O    0.0    0.2    1.0    CX1-
CX2->C14-C12,
Network11  bmse000983 O_acetyl_L_serine  D2O    1.0    0.333  1.0    CX1-
CX2->C10-C9,
Network12  bmse000019 D_ornithine    D2O    0.0    0.25   1.0    CX1-CX2-
>C4-C5,
Network12  bmse000109 putrescine D2O    0.0    0.333  1.0    CX1-CX2->C5-
C4,
Network12  bmse000116 spermidine D2O    1.0    0.2    1.0    CX1-CX2->C8-
C9,
Network12  bmse000162 L_ornithine    D2O    0.0    0.25   1.0    CX1-CX2-
>C4-C5,
Network12  bmse000195 N_alpha_acetyl_DL_ornithine    D2O    0.0    0.333
1.0    CX1-CX2->C8-C11,
Network12  bmse000249 methyl_4_aminobutyrate D2O    0.0    0.333  1.0
CX1-CX2->C2-C1,
Network12  bmse000340 gamma_Aminobutyric_acid    D2O    0.0    0.333  1.0
CX1-CX2->C4-C6,
Network12  bmse000468 4_Acetamidobutyric_acid    D2O    0.0    0.25   1.0
CX1-CX2->C5-C6,
Network12  bmse000862 putrescine D2O    0.0    0.333  1.0    CX1-CX2->C5-
C4,
```

```
Network12  bmse000871 gamma_Aminobutyric_acid    D2O    0.0    0.333  1.0
CX1-CX2->C4-C6,
Network12  bmse000897 D_ornithine    D2O    0.0    0.25   1.0    CX1-CX2-
>C4-C5,
Network12  bmse000951 spermidine D2O    1.0    0.2    1.0    CX1-CX2->C8-
C9,
Network15  bmse000047 L_proline  D2O    0.0    0.25   1.0    CX2-CX1->C6-
C7,
Network15  bmse000116 spermidine D2O    1.0    0.2    1.0    CX2-CX1->C8-
C7,
Network15  bmse000947 L_proline  D2O    0.0    0.25   1.0    CX2-CX1->C6-
C7,
Network15  bmse000951 spermidine D2O    1.0    0.2    1.0    CX2-CX1->C8-
C7,
Network17  bmse000037 L_glutamic_acid    D2O    0.0    0.25   1.0    CX2-
CX1->C4-C6,
Network17  bmse000038 L_glutamine    D2O    0.0    0.25   1.0    CX2-CX1-
>C2-C5,
Network17  bmse000194 L_arginine_L_glutamate D2O    0.0    0.2    1.0
CX2-CX1->C4-C2,
Network18  bmse000038 L_glutamine    D2O    0.0    0.5    0.667  CX5-CX3-
>C6-C5,CX5-CX2->C6-C6,CX4-CX2->C7-C6,  CX1-CX3->?-?,CX1-CX6->?-?,
Network18  bmse000565 cis_5_dodecenoic_acid  CDCl3  0.0    0.273  0.667
CX5-CX3->C3-C5,CX5-CX2->C3-C13,CX4-CX2->C14-C13,   CX1-CX3->?-?,CX1-CX6-
>?-?,
Network18  bmse000625 4_methylvaleric_acid   CDCl3  0.0    0.4    0.5
CX5-CX2->C4-C5,CX4-CX2->C8-C5, CX1-CX3->?-?,CX1-CX6->?-?,CX5-CX3->C4-?,
Network19  bmse000019 D_ornithine    D2O    0.0    0.5    0.8    CX4-CX2-
>C2-C3,CX4-CX5->C2-C2,CX5-CX5->C2-C2,CX5-CX1->C2-C1,   CX3-CX5->?-?,
Network19  bmse000029 L_arginine D2O    0.0    0.5    0.8    CX4-CX2->C2-
C5,CX4-CX5->C2-C2,CX5-CX5->C2-C2,CX5-CX1->C2-C3,   CX3-CX5->?-?,
Network19  bmse000032 L_citrulline   D2O    0.0    0.5    0.8    CX4-CX2-
>C4-C6,CX4-CX5->C4-C4,CX5-CX5->C4-C4,CX5-CX1->C4-C1,   CX3-CX5->?-?,
Network19  bmse000037 L_glutamic_acid    D2O    0.0    0.5    0.8    CX4-
CX2->C4-C6,CX4-CX5->C4-C4,CX5-CX5->C4-C4,CX5-CX1->C4-C1,   CX3-CX5->?-?,
Network19  bmse000038 L_glutamine    D2O    0.0    0.5    0.8    CX4-CX2-
>C2-C5,CX4-CX5->C2-C2,CX5-CX5->C2-C2,CX5-CX1->C2-C3,   CX3-CX5->?-?,
Network19  bmse000039 L_histidine    D2O    0.333  0.5    0.8    CX4-CX2-
>C5-C3,CX4-CX5->C5-C5,CX5-CX5->C5-C5,CX5-CX1->C5-C6,   CX3-CX5->?-?,
Network19  bmse000162 L_ornithine    D2O    0.0    0.5    0.8    CX4-CX2-
>C2-C3,CX4-CX5->C2-C2,CX5-CX5->C2-C2,CX5-CX1->C2-C7,   CX3-CX5->?-?,
Network19  bmse000185 L_glutathione_reduced  D2O    0.0    0.429  1.0
CX4-CX2->C2-C6,CX4-CX5->C2-C2,CX5-CX5->C2-C2,CX5-CX1->C2-C3,CX3-CX5->C12-
C2,
Network19  bmse000209 D_citrulline   D2O    0.0    0.5    0.8    CX4-CX2-
>C6-C11,CX4-CX5->C6-C6,CX5-CX5->C6-C6,CX5-CX1->C6-C2,  CX3-CX5->?-?,
Network19  bmse000289 S_Adenosyl_L_homocysteine  D2O    0.143  0.222  0.8
CX4-CX2->C13-C15,CX4-CX5->C13-C13,CX5-CX5->C13-C13,CX5-CX1->C13-C21,
CX3-CX5->?-?,
Network19  bmse000449 Npai_Methyl_L_histidine    D2O    0.0    0.25   0.6
CX4-CX5->C7-C7,CX5-CX5->C7-C7,CX5-CX1->C7-C11, CX4-CX2->C7-?,CX3-CX5->?-?,
Network19  bmse000711 L_arginine D2O    0.0    0.5    0.8    CX4-CX2->C2-
C5,CX4-CX5->C2-C2,CX5-CX5->C2-C2,CX5-CX1->C2-C3,   CX3-CX5->?-?,
Network19  bmse000858 L_citrulline   D2O    0.0    0.5    0.8    CX4-CX2-
```

```
>C4-C6,CX4-CX5->C4-C4,CX5-CX5->C4-C4,CX5-CX1->C4-C1,   CX3-CX5->?-?,
Network19  bmse000897 D_ornithine    D2O    0.0    0.5    0.8    CX4-CX2-
>C2-C3,CX4-CX5->C2-C2,CX5-CX5->C2-C2,CX5-CX1->C2-C1,   CX3-CX5->?-?,
Network19  bmse000952 L_glutathione_reduced  D2O    0.0    0.429  1.0
CX4-CX2->C2-C6,CX4-CX5->C2-C2,CX5-CX5->C2-C2,CX5-CX1->C2-C3,CX3-CX5->C12-
C2,
Network21  bmse000483 R_2_Pyrrolidinone_5_carboxylate    D2O    0.0
0.25   1.0    CX1-CX2->C7-C8,
Network21  bmse000878 R_2_Pyrrolidinone_5_carboxylate    D2O    0.0
0.25   1.0    CX1-CX2->C7-C8,
Network22  bmse000033 L_cystathionine    D2O    0.0    0.333  1.0    CX1-
CX2->C2-C10,
Network22  bmse000043 L_lysine   D2O    0.0    0.2    1.0    CX1-CX2->C5-
C2,
Network22  bmse000044 L_methionine   D2O    0.0    0.333  1.0    CX1-CX2-
>C5-C2,
Network22  bmse000169 L_selenomethionine D2O    0.0    0.333  1.0    CX1-
CX2->C6-C2,
Network22  bmse000291 Selenomethionine   D2O    0.0    0.333  1.0    CX1-
CX2->C7-C5,
Network22  bmse000411 L_Norleucine   D2O    0.0    0.2    1.0    CX1-CX2-
>C4-C6,
Network22  bmse000429 DL_2_Aminoadipic_acid  D2O    0.0    0.2    1.0
CX1-CX2->C6-C8,
Network22  bmse000442 L_homocitrulline   D2O    0.0    0.2    1.0    CX1-
CX2->C8-C10,
Network22  bmse000450 D_Methionine   D2O    0.0    0.333  1.0    CX1-CX2-
>C5-C6,
Network22  bmse000466 DL_beta_leucine    D2O    0.0    0.2    1.0    CX1-
CX2->C4-C5,
Network22  bmse000745 homoarginine   D2O    0.0    0.2    1.0    CX1-CX2-
>C8-C10,
Network23  bmse000493 epsilon_caprolactone   CDCl3  0.0    0.2    1.0
CX1-CX2->C8-C6,
Network24  bmse000159 beta_alanine   D2O    0.0    0.5    0.5    CX4-CX1-
>C2-C6,    CX1-CX3->C6-?,CX3-CX2->?-?,
Network24  bmse000388 Cysteamine D2O    0.0    1.0    0.5    CX4-CX1->C4-
C3,    CX1-CX3->C3-?,CX3-CX2->?-?,
Network24  bmse000967 beta_alanine   D2O    0.0    0.5    0.5    CX4-CX1-
>C2-C6,    CX1-CX3->C6-?,CX3-CX2->?-?,
Network25  bmse000452 Hypotaurine    D2O    0.0    1.0    1.0    CX1-CX2-
>C5-C6,
Network26  bmse000037 L_glutamic_acid    D2O    0.0    0.25   1.0    CX1-
CX2->C2-C7,
Network26  bmse000794 3_carboxypropyl_trimethyl_ammonium D2O    0.571
0.333  1.0    CX1-CX2->C10-C9,
Network27  bmse000037 L_glutamic_acid    D2O    0.0    0.25   1.0    CX1-
CX2->C7-C2,
Network27  bmse000183 succinic_acid  D2O    0.0    0.333  1.0    CX1-CX2-
>C6-C2,
Network27  bmse000340 gamma_Aminobutyric_acid    D2O    0.0    0.333  1.0
CX1-CX2->C5-C7,
Network27  bmse000344 4_Guanidinobutyric_acid    D2O    0.0    0.333  1.0
CX1-CX2->C7-C9,
```

```
Network27  bmse000376 N_carbamyl_L_glutamate D2O    0.0    0.25   1.0
CX1-CX2->C10-C12,
Network27  bmse000382 N_Acetyl_L_glutamic_acid   D2O    0.0    0.2    1.0
CX1-CX2->C9-C12,
Network27  bmse000468 4_Acetamidobutyric_acid    D2O    0.0    0.25   1.0
CX1-CX2->C7-C8,
Network27  bmse000871 gamma_Aminobutyric_acid    D2O    0.0    0.333  1.0
CX1-CX2->C5-C7,
Network29  bmse000030 L_asparagine   D2O    0.5    0.333  0.5    CX2-CX1-
>C6-C5,    CX2-CX3->C6-?,CX3-CX4->?-?,
Network29  bmse000041 L_isoleucine   D2O    0.0    0.2    0.5    CX3-CX4-
>C4-C3,    CX2-CX1->?-?,CX2-CX3->?-?,
Network29  bmse000049 L_threonine    D2O    0.0    0.333  0.5    CX3-CX4-
>C4-C3,    CX2-CX1->?-?,CX2-CX3->?-?,
Network29  bmse000052 L_valine   D2O    0.4    0.25   0.5    CX3-CX4->C3-
C2,    CX2-CX1->?-?,CX2-CX3->?-?,
Network29  bmse000123 trans_4_hydroxy_L_proline  D2O    0.0    0.25   0.5
CX3-CX4->C4-C1,    CX2-CX1->?-?,CX2-CX3->?-?,
Network29  bmse000148 diethyl_oxalacetate    D2O    1.0    0.2    0.5
CX3-CX4->C7-C6,    CX2-CX1->?-?,CX2-CX3->?-?,
Network29  bmse000631 O_phospho_DL_threonine D2O    0.0    0.333  0.5
CX3-CX4->C12-C10,  CX2-CX1->?-?,CX2-CX3->?-?,
Network29  bmse000741 N_methyl_L_aspartic_acid   D2O    0.4    1.0    1.0
CX2-CX1->C10-C7,CX2-CX3->C10-C8,CX3-CX4->C8-C6,
Network29  bmse000859 L_threonine    D2O    0.0    0.333  0.5    CX3-CX4-
>C4-C3,    CX2-CX1->?-?,CX2-CX3->?-?,
Network29  bmse000860 L_valine   D2O    0.4    0.25   0.5    CX3-CX4->C3-
C2,    CX2-CX1->?-?,CX2-CX3->?-?,
Network29  bmse000866 L_isoleucine   D2O    0.0    0.2    0.5    CX3-CX4-
>C4-C3,    CX2-CX1->?-?,CX2-CX3->?-?,
Network29  bmse000966 trans_4_hydroxy_L_proline  D2O    0.0    0.25   0.5
CX3-CX4->C4-C1,    CX2-CX1->?-?,CX2-CX3->?-?,
Network30  bmse000120 taurine    D2O    0.0    1.0    1.0    CX1-CX2->C2-
C7,
Network31  bmse000042 L_leucine  D2O    0.333  0.2    1.0    CX1-CX2->C5-
C2,
Network31  bmse000423 N_acetyl_L_aspartic_acid   D2O    0.0    0.25   1.0
CX1-CX2->C8-C7,
Network32  bmse000042 L_leucine  D2O    0.333  0.2    1.0    CX2-CX1->C5-
C2,
Network32  bmse000423 N_acetyl_L_aspartic_acid   D2O    0.0    0.25   1.0
CX2-CX1->C8-C7,
Network32  bmse000453 Ureidosuccinic_acid    D2O    0.0    0.333  1.0
CX2-CX1->C9-C8,
Network33  bmse000308 2_Aminoethyl_dihydrogen_phosphate  D2O    0.0    1.0
1.0    CX2-CX1->C7-C8,
Network36  bmse000276 Ethanolamine   D2O    0.0    1.0    1.0    CX2-CX1-
>C4-C3,
Network37  bmse000074 D_carnitine    D2O    0.571  0.333  1.0    CX1-CX2-
>C9-C8,
Network37  bmse000211 L_carnitine    D2O    0.0    0.333  1.0    CX1-CX2-
>C1-C2,
Network37  bmse000949 D_carnitine    D2O    0.571  0.333  1.0    CX1-CX2-
>C9-C8,
```

```
Network38  bmse000367 chloroacetic_acid  D2O    0.0    1.0    0.5    CX3-
CX3->C4-C4,CX1-CX3->C5-C4,CX1-CX1->C5-C5,  CX3-CX2->C4-?,CX4-CX4->?-?,CX2-
CX4->?-?,CX2-CX2->?-?,CX2-CX1->?-?,CX1-CX4->C5-?,
Network38  bmse000658 phenylacetylglycine    D2O    0.0    0.333  0.75
CX3-CX3->C10-C10,CX3-CX2->C10-C14,CX2-CX2->C14-C14,CX2-CX1->C14-C7,CX1-
CX3->C7-C10,CX1-CX1->C7-C7, CX4-CX4->?-?,CX2-CX4->C14-?,CX1-CX4->C7-?,
Network41  bmse000028 L_alanine  D2O    0.0    0.5    1.0    CX1-CX2->C2-
C3,
Network41  bmse000236 D_alanine  D2O    0.0    0.5    1.0    CX1-CX2->C2-
C4,
Network41  bmse000282 DL_Alanine D2O    0.0    0.5    1.0    CX1-CX2->C4-
C6,
Network43  bmse000019 D_ornithine    D2O    0.0    0.25   1.0    CX1-CX2-
>C1-C2,
Network43  bmse000038 L_glutamine    D2O    0.0    0.25   1.0    CX1-CX2-
>C3-C2,
Network43  bmse000040 L_homoserine   D2O    0.0    0.333  1.0    CX1-CX2-
>C3-C2,
Network43  bmse000042 L_leucine  D2O    0.333  0.2    1.0    CX1-CX2->C3-
C2,
Network43  bmse000044 L_methionine   D2O    0.0    0.333  1.0    CX1-CX2-
>C3-C2,
Network43  bmse000073 L_canavanine   D2O    0.0    0.333  1.0    CX1-CX2-
>C5-C2,
Network43  bmse000078 creatine   D2O    0.0    1.0    1.0    CX1-CX2->C4-
C5,
Network43  bmse000162 L_ornithine    D2O    0.0    0.25   1.0    CX1-CX2-
>C7-C2,
Network43  bmse000430 DL_homocysteine    D2O    0.0    0.333  1.0    CX1-
CX2->C8-C5,
Network43  bmse000450 D_Methionine   D2O    0.0    0.333  1.0    CX1-CX2-
>C8-C6,
Network43  bmse000897 D_ornithine    D2O    0.0    0.25   1.0    CX1-CX2-
>C1-C2,
Network43  bmse000950 creatine   D2O    0.0    1.0    1.0    CX1-CX2->C4-
C5,
Network43  bmse000983 O_acetyl_L_serine  D2O    1.0    0.333  1.0    CX1-
CX2->C8-C6,
Network44  bmse000285 Choline    D2O    0.8    1.0    1.0    CX1-CX2->C3-
C4,
Network44  bmse000953 Choline    D2O    0.8    1.0    1.0    CX1-CX2->C3-
C4,
Network47  bmse000049 L_threonine    D2O    0.0    0.333  1.0    CX2-CX1-
>C3-C4,
Network47  bmse000148 diethyl_oxalacetate    D2O    1.0    0.2    1.0
CX2-CX1->C6-C7,
Network47  bmse000631 O_phospho_DL_threonine D2O    0.0    0.333  1.0
CX2-CX1->C10-C12,
Network47  bmse000859 L_threonine    D2O    0.0    0.333  1.0    CX2-CX1-
>C3-C4,
Network48  bmse000008 D_allose   D2O    0.667  0.4    0.75   CX4-CX3->C4-
C1,CX1-CX3->C9-C1, CX2-CX1->?-?,
Network48  bmse000013 D_galactose    D2O    1.0    0.2    0.75   CX4-CX3-
>C6-C5,CX1-CX3->C5-C5, CX2-CX1->?-?,
```

```
Network48  bmse000017 D_maltose  D2O    0.917  0.2    0.75   CX4-CX3->C6-
C2,CX1-CX3->C8-C2, CX2-CX1->?-?,
Network48  bmse000018 D_mannose  D2O    1.0    0.2    0.75   CX4-CX3->C7-
C6,CX1-CX3->C6-C6, CX2-CX1->?-?,
Network48  bmse000022 D_sorbose  D2O    0.5    0.4    0.75   CX4-CX3->C8-
C10,CX1-CX3->C6-C10,   CX2-CX1->?-?,
Network48  bmse000023 D_tagatose D2O    0.833  0.4    0.75   CX4-CX3->C8-
C10,CX1-CX3->C6-C10,   CX2-CX1->?-?,
Network48  bmse000026 D_xylose   D2O    1.0    0.25   0.75   CX4-CX3->C5-
C4,CX1-CX3->C4-C4, CX2-CX1->?-?,
Network48  bmse000086 alpha_D_glucose_1_phosphate    D2O    0.0    0.4
1.0    CX4-CX3->C6-C5,CX2-CX1->C4-C5,CX1-CX3->C5-C5,
Network48  bmse000087 alpha_D_glucose_1_6_bisphosphate   D2O    0.0    0.2
0.5    CX2-CX1->C4-C5,    CX4-CX3->?-?,CX1-CX3->C5-?,
Network48  bmse000099 D_mannitol D2O    0.0    0.4    0.75   CX2-CX1->C7-
C9,CX1-CX3->C9-C3, CX4-CX3->?-?,
Network48  bmse000115 D_sorbitol D2O    1.0    0.2    0.75   CX4-CX3->C6-
C5,CX1-CX3->C5-C5, CX2-CX1->?-?,
Network48  bmse000138 D_cellobiose   D2O    1.0    0.2    0.75   CX4-CX3-
>C6-C2,CX1-CX3->C8-C2, CX2-CX1->?-?,
Network48  bmse000151 alpha_D_galactose_1_phosphate  D2O    0.0    0.4
1.0    CX4-CX3->C6-C5,CX2-CX1->C4-C5,CX1-CX3->C5-C5,
Network48  bmse000163 N_acetyl_D_glucosamine_1_phosphate D2O    0.625
0.333  0.75   CX4-CX3->C15-C10,CX1-CX3->C7-C10,  CX2-CX1->?-?,
Network48  bmse000189 D_glucosamine_6_phosphate  D2O    0.0    0.2    0.5
CX2-CX1->C3-C4,    CX4-CX3->?-?,CX1-CX3->C4-?,
Network48  bmse000233 melibiose  D2O    0.0    0.2    0.5    CX2-CX1->C9-
C7,    CX4-CX3->?-?,CX1-CX3->C7-?,
Network48  bmse000235 L_gulonolactone    D2O    0.0    0.6    1.0    CX4-
CX3->C8-C4,CX2-CX1->C2-C5,CX1-CX3->C5-C4,
Network48  bmse000865 D_tagatose D2O    0.833  0.4    0.75   CX4-CX3->C8-
C10,CX1-CX3->C6-C10,   CX2-CX1->?-?,
Network48  bmse000898 D_xylose   D2O    1.0    0.25   0.75   CX4-CX3->C5-
C4,CX1-CX3->C4-C4, CX2-CX1->?-?,
Network48  bmse000903 D_sorbose  D2O    0.5    0.4    0.75   CX4-CX3->C8-
C10,CX1-CX3->C6-C10,   CX2-CX1->?-?,
Network48  bmse000939 D_cellobiose   D2O    1.0    0.2    0.75   CX4-CX3-
>C6-C2,CX1-CX3->C8-C2, CX2-CX1->?-?,
Network48  bmse000946 D_maltose  D2O    0.917  0.2    0.75   CX4-CX3->C6-
C2,CX1-CX3->C8-C2, CX2-CX1->?-?,
Network48  bmse001006 D_galactose    D2O    1.0    0.2    0.75   CX4-CX3-
>C6-C5,CX1-CX3->C5-C5, CX2-CX1->?-?,
Network48  bmse001007 D_sorbitol D2O    1.0    0.2    0.75   CX4-CX3->C6-
C5,CX1-CX3->C5-C5, CX2-CX1->?-?,
Network48  bmse001008 D_allose   D2O    0.667  0.4    0.75   CX4-CX3->C4-
C1,CX1-CX3->C9-C1, CX2-CX1->?-?,
Network50  bmse000008 D_allose   D2O    0.667  0.2    1.0    CX2-CX1->C4-
C1,
Network50  bmse000010 D_fructose D2O    1.0    0.2    1.0    CX2-CX1->C11-
C8,
Network50  bmse000013 D_galactose    D2O    1.0    0.2    1.0    CX2-CX1-
>C6-C5,
Network50  bmse000018 D_mannose  D2O    1.0    0.2    1.0    CX2-CX1->C7-
C6,
```

```
Network50  bmse000022 D_sorbose  D2O    0.5    0.2    1.0    CX2-CX1->C8-
C10,
Network50  bmse000023 D_tagatose D2O    0.833  0.2    1.0    CX2-CX1->C8-
C10,
Network50  bmse000062 adonitol   D2O    0.0    0.25   1.0    CX2-CX1->C9-
C7,
Network50  bmse000068 L_arabitol D2O    0.0    0.25   1.0    CX2-CX1->C7-
C3,
Network50  bmse000084 gluconic_acid  D2O    0.0    1.0    1.0    CX2-CX1-
>C6-C5,
Network50  bmse000087 alpha_D_glucose_1_6_bisphosphate   D2O    0.0    0.2
1.0    CX2-CX1->C6-C5,
Network50  bmse000095 i_erythritol   D2O    0.0    0.333  1.0    CX2-CX1-
>C5-C4,
Network50  bmse000099 D_mannitol D2O    0.0    0.2    1.0    CX2-CX1->C2-
C3,
Network50  bmse000100 meso_erythritol    D2O    0.0    0.333  1.0    CX2-
CX1->C5-C4,
Network50  bmse000115 D_sorbitol D2O    1.0    0.2    1.0    CX2-CX1->C6-
C5,
Network50  bmse000121 L_threitol D2O    0.0    0.333  1.0    CX2-CX1->C5-
C4,
Network50  bmse000129 xylitol    D2O    0.0    0.25   1.0    CX2-CX1->C5-
C4,
Network50  bmse000151 alpha_D_galactose_1_phosphate  D2O    0.0    0.2
1.0    CX2-CX1->C6-C5,
Network50  bmse000184 glycerol   D2O    0.0    0.5    1.0    CX2-CX1->C6-
C1,
Network50  bmse000193 DL_alpha_glycerol_phosphate    D2O    0.0    0.5
1.0    CX2-CX1->C10-C7,
Network50  bmse000230 D_glucono_1_5_lactone  D2O    0.0    1.0    1.0
CX2-CX1->C6-C5,
Network50  bmse000235 L_gulonolactone    D2O    0.0    0.2    1.0    CX2-
CX1->C8-C4,
Network50  bmse000440 D_Glucosaminic_acid    D2O    0.0    0.2    1.0
CX2-CX1->C12-C10,
Network50  bmse000856 glycerol   D2O    0.0    0.5    1.0    CX2-CX1->C6-
C1,
Network50  bmse000865 D_tagatose D2O    0.833  0.2    1.0    CX2-CX1->C8-
C10,
Network50  bmse000869 L_arabitol D2O    0.0    0.25   1.0    CX2-CX1->C7-
C3,
Network50  bmse000903 D_sorbose  D2O    0.5    0.2    1.0    CX2-CX1->C8-
C10,
Network50  bmse001005 D_fructose D2O    1.0    0.2    1.0    CX2-CX1->C11-
C8,
Network50  bmse001006 D_galactose    D2O    1.0    0.2    1.0    CX2-CX1-
>C6-C5,
Network50  bmse001007 D_sorbitol D2O    1.0    0.2    1.0    CX2-CX1->C6-
C5,
Network50  bmse001008 D_allose   D2O    0.667  0.2    1.0    CX2-CX1->C4-
C1,
Network52  bmse000204 D_ribose_5_phosphate   D2O    1.0    0.25   1.0
CX1-CX2->C7-C6,
```

```
Network53  bmse000018 D_mannose  D2O    1.0    0.2    1.0    CX2-CX1->C7-
C6,
Network53  bmse000193 DL_alpha_glycerol_phosphate    D2O    0.0    0.5
1.0    CX2-CX1->C4-C7,
Network53  bmse000257 FMN    D2O    0.0    1.0    1.0    CX2-CX1->C18-C16,
Network54  bmse000069 betaine    D2O    0.8    1.0    1.0    CX1-CX2->C4-
C3,
Network54  bmse000948 betaine    D2O    0.8    1.0    1.0    CX1-CX2->C4-
C3,
Network55  bmse000208 L_lactic_acid  D2O    0.0    0.5    1.0    CX2-CX1-
>C1-C2,
Network55  bmse000269 R_Lactate  D2O    0.0    0.5    1.0    CX2-CX1->C6-
C4,
Network55  bmse000979 L_lactic_acid  D2O    0.0    0.5    1.0    CX2-CX1-
>C1-C2,
Network56  bmse000008 D_allose   D2O    0.667  0.2    0.75   CX4-CX2->C9-
C9,CX4-CX1->C9-C7, CX2-CX3->C9-?,
Network56  bmse000013 D_galactose    D2O    1.0    0.4    0.75   CX4-CX2-
>C2-C5,CX4-CX1->C2-C1, CX2-CX3->C5-?,
Network56  bmse000015 D_glucose  D2O    1.0    0.2    0.75   CX4-CX1->C2-
C1,    CX4-CX2->C2-?,CX2-CX3->?-?,
Network56  bmse000018 D_mannose  D2O    1.0    0.4    0.75   CX4-CX2->C3-
C6,CX4-CX1->C3-C2, CX2-CX3->C6-?,
Network56  bmse000026 D_xylose   D2O    1.0    0.25   0.75   CX4-CX1->C2-
C1,    CX4-CX2->C2-?,CX2-CX3->?-?,
Network56  bmse000086 alpha_D_glucose_1_phosphate    D2O    0.0    0.2
0.75   CX4-CX2->C2-C3,CX2-CX3->C3-C3, CX4-CX1->C2-?,
Network56  bmse000125 D_trehalose    D2O    0.833  0.2    0.75   CX4-CX2-
>C14-C5,CX4-CX1->C14-C13,  CX2-CX3->C5-?,
Network56  bmse000233 melibiose  D2O    0.0    0.2    0.5    CX2-CX3->C20-
C17,  CX4-CX2->?-?,CX4-CX1->?-?,
Network56  bmse000239 D_saccharate   D2O    0.0    0.2    0.5    CX2-CX3-
>C8-C6,    CX4-CX2->?-?,CX4-CX1->?-?,
Network56  bmse000855 D_glucose  D2O    1.0    0.2    0.75   CX4-CX1->C2-
C1,    CX4-CX2->C2-?,CX2-CX3->?-?,
Network56  bmse000876 D_trehalose    D2O    0.833  0.2    0.75   CX4-CX2-
>C14-C5,CX4-CX1->C14-C13,  CX2-CX3->C5-?,
Network56  bmse000898 D_xylose   D2O    1.0    0.25   0.75   CX4-CX1->C2-
C1,    CX4-CX2->C2-?,CX2-CX3->?-?,
Network56  bmse001006 D_galactose    D2O    1.0    0.4    0.75   CX4-CX2-
>C2-C5,CX4-CX1->C2-C1, CX2-CX3->C5-?,
Network56  bmse001008 D_allose   D2O    0.667  0.2    0.75   CX4-CX2->C9-
C9,CX4-CX1->C9-C7, CX2-CX3->C9-?,
Network57  bmse000140 D_glucuronate  D2O    0.0    1.0    1.0    CX3-CX4-
>C2-C2,CX1-CX3->C1-C2,CX2-CX4->C3-C2,
Network57  bmse000313 beta_gentiobiose   D2O    0.0    0.2    1.0    CX3-
CX4->C20-C20,CX1-CX3->C18-C20,CX2-CX4->C22-C20,
Network59  bmse000346 Barbituric_acid    D2O    0.0    0.5    1.0    CX1-
CX2->C6-C8,
Network63  bmse000039 L_histidine    D2O    0.333  0.25   1.0    CX1-CX2-
>C1-C4,
Network63  bmse000077 coumarin   CDCl3  0.0    0.25   1.0    CX1-CX2->C4-
C2,
Network63  bmse000246 L_carnosine    D2O    0.0    0.25   1.0    CX1-CX2-
```

```
>C1-C6,
Network63  bmse000448 L_Histidinol   D2O    0.0    0.25   1.0    CX1-CX2-
>C9-C7,
Network63  bmse000707 N_acetyl_histidine D2O    0.0    0.2    1.0    CX1-
CX2->C11-C9,
Network63  bmse000744 histamine  D2O    0.0    0.333  1.0    CX1-CX2->C7-
C5,
Network63  bmse000762 neostigmine    D2O    0.167  0.25   1.0    CX1-CX2-
>C11-C13,
Network65  bmse000281 Nicotinamide   D2O    0.0    0.2    1.0    CX1-CX2-
>C9-C7,
Network65  bmse000622 3_pyridinecarbonitrile D2O    0.0    0.2    1.0
CX1-CX2->C5-C4,
Network66  bmse000432 Pyridine   D2O    0.0    0.25   1.0    CX1-CX2->C6-
C4,
```

Supplementary Information 5: Example of MatchJRES in PyINETA.

```
Network1
"/Users/rtaujale/Dropbox/Projects/collaborative/13C_mouse_INETA/raw
_data/pyINETA_AlaRef_2023/3.tilt.sym.ft2=CX1[Present](27.03,40.91):1129341
600000.0;CX2[Present](13.72,41.01):688701440000.0;"

Network2
"/Users/rtaujale/Dropbox/Projects/collaborative/13C_mouse_INETA/raw_data/p
yINETA_AlaRef_2023/3.tilt.sym.ft2=CX1[Present](38.58,56.17):772994430000.0
;CX2[Present](17.27,55.93):827546400000.0;"

Network3
"/Users/rtaujale/Dropbox/Projects/collaborative/13C_mouse_INETA/raw_data/p
yINETA_AlaRef_2023/3.tilt.sym.ft2=CX1[Present](18.7,72.18):2646701000000.0
;CX2[Present](53.2,71.98):1644530800000.0;"

Network4
"/Users/rtaujale/Dropbox/Projects/collaborative/13C_mouse_INETA/raw_data/p
yINETA_AlaRef_2023/3.tilt.sym.ft2=CX1[Present](31.73,51.09):1181092500000.
0;CX2[Present](19.2,50.97):1298547300000.0;"

Network5
"/Users/rtaujale/Dropbox/Projects/collaborative/13C_mouse_INETA/raw_data/p
yINETA_AlaRef_2023/3.tilt.sym.ft2=CX1[Present](31.82,52.36):1180879700000.
0;CX2[Present](20.57,52.45):1632450800000.0;"

Network6
"/Users/rtaujale/Dropbox/Projects/collaborative/13C_mouse_INETA/raw_data/p
yINETA_AlaRef_2023/3.tilt.sym.ft2=CX1[Present](68.52,90.83):1235223000000.
0;CX2[Present](22.24,90.78):832188700000.0;"

Network7
"/Users/rtaujale/Dropbox/Projects/collaborative/13C_mouse_INETA/raw_data/p
yINETA_AlaRef_2023/3.tilt.sym.ft2=CX1[Present](22.87,94.29):5607700600000.
0;CX3[Present](70.98,94.12):2692579000000.0;"

Network8
"/Users/rtaujale/Dropbox/Projects/collaborative/13C_mouse_INETA/raw_data/p
yINETA_AlaRef_2023/3.tilt.sym.ft2=CX1[Present](24.7,51.54):1726148600000.0
;CX2[Present](42.71,69.7):1706099900000.0;CX4[Present](26.95,69.7):1141805
800000.0;CX5[Present](64.76,107.56):1144215200000.0;CX8[Present](23.43,50.
35):1606211000000.0;"

Network9
"/Users/rtaujale/Dropbox/Projects/collaborative/13C_mouse_INETA/raw_data/p
yINETA_AlaRef_2023/3.tilt.sym.ft2=CX1[Present](24.13,53.52):815090300000.0
;CX2[Present](29.14,53.52):4477700000000.0;"

Network10
"/Users/rtaujale/Dropbox/Projects/collaborative/13C_mouse_INETA/raw_data/p
yINETA_AlaRef_2023/3.tilt.sym.ft2=CX1[Present](32.59,57.04):1074266700000.
0;CX2[Present](24.3,56.98):834897440000.0;"
Network11
```

```
"/Users/rtaujale/Dropbox/Projects/collaborative/13C_mouse_INETA/raw_data/p
yINETA_AlaRef_2023/3.tilt.sym.ft2=CX1[Present](24.64,201.46):1738039100000
.0;CX2[Present](176.75,201.46):912965600000.0;"

Network12
"/Users/rtaujale/Dropbox/Projects/collaborative/13C_mouse_INETA/raw_data/p
yINETA_AlaRef_2023/3.tilt.sym.ft2=CX1[Present](25.69,67.17):993319650000.0
;CX2[Present](41.51,67.17):1120700700000.0;"

Network13
"/Users/rtaujale/Dropbox/Projects/collaborative/13C_mouse_INETA/raw_data/p
yINETA_AlaRef_2023/3.tilt.sym.ft2=CX1[Present](183.03,208.87):779128500000
.0;CX2[Present](25.8,209.03):980739600000.0;"

Network14
"/Users/rtaujale/Dropbox/Projects/collaborative/13C_mouse_INETA/raw_data/p
yINETA_AlaRef_2023/3.tilt.sym.ft2=CX1[Present](26.42,86.41):766211300000.0
;CX2[Present](60.42,86.62):740303960000.0;"

Network15
"/Users/rtaujale/Dropbox/Projects/collaborative/13C_mouse_INETA/raw_data/p
yINETA_AlaRef_2023/3.tilt.sym.ft2=CX1[Present](48.6,75.22):741234100000.0;
CX2[Present](26.53,75.15):762627160000.0;"

Network16
"/Users/rtaujale/Dropbox/Projects/collaborative/13C_mouse_INETA/raw_data/p
yINETA_AlaRef_2023/3.tilt.sym.ft2=CX1[Present](182.24,243.22):820035850000
.0;CX2[Present](28.07,89.06):1381270400000.0;CX3[Present](60.97,243.13):10
34876350000.0;"

Network17
"/Users/rtaujale/Dropbox/Projects/collaborative/13C_mouse_INETA/raw_data/p
yINETA_AlaRef_2023/3.tilt.sym.ft2=CX1[Present](28.95,85.85):4472378500000.
0;CX2[Present](56.68,85.82):7125908600000.0;"

Network18
"/Users/rtaujale/Dropbox/Projects/collaborative/13C_mouse_INETA/raw_data/p
yINETA_AlaRef_2023/3.tilt.sym.ft2=CX1[Present](29.03,70.84):4478046600000.
0;CX2[Present](33.61,213.39):2013865600000.0;CX4[Present](179.7,213.2):931
228200000.0;CX6[Present](41.66,71.0):1278682200000.0;"

Network19
"/Users/rtaujale/Dropbox/Projects/collaborative/13C_mouse_INETA/raw_data/p
yINETA_AlaRef_2023/3.tilt.sym.ft2=CX1[Present](176.46,233.69):876906100000
.0;CX2[Present](29.51,86.95):1781145800000.0;CX4[Present](57.33,86.94):129
4262500000.0;"

Network20
"/Users/rtaujale/Dropbox/Projects/collaborative/13C_mouse_INETA/raw_data/p
yINETA_AlaRef_2023/3.tilt.sym.ft2=CX1[Present](32.53,64.2):1074889560000.0
;"

Network21
```

```
"/Users/rtaujale/Dropbox/Projects/collaborative/13C_mouse_INETA/raw_data/p
yINETA_AlaRef_2023/3.tilt.sym.ft2=CX1[Present](32.4,216.14):997792700000.0
;CX2[Present](183.72,216.19):773462160000.0;"

Network22
"/Users/rtaujale/Dropbox/Projects/collaborative/13C_mouse_INETA/raw_data/p
yINETA_AlaRef_2023/3.tilt.sym.ft2=CX1[Present](32.76,89.8):863568660000.0;
CX2[Present](57.01,89.9):3025168000000.0;"

Network23
"/Users/rtaujale/Dropbox/Projects/collaborative/13C_mouse_INETA/raw_data/p
yINETA_AlaRef_2023/3.tilt.sym.ft2=CX1[Present](176.93,210.88):917193600000
.0;CX2[Present](34.1,210.98):1200629400000.0;"

Network24
"/Users/rtaujale/Dropbox/Projects/collaborative/13C_mouse_INETA/raw_data/p
yINETA_AlaRef_2023/3.tilt.sym.ft2=CX1[Present](39.26,75.41):1042545840000.
0;CX2[Present](40.64,80.07):698567500000.0;CX4[Present](35.96,75.41):10097
21000000.0;"

Network25
"/Users/rtaujale/Dropbox/Projects/collaborative/13C_mouse_INETA/raw_data/p
yINETA_AlaRef_2023/3.tilt.sym.ft2=CX1[Present](57.67,93.92):1002780160000.
0;CX2[Present](36.09,94.03):1026640250000.0;"

Network26
"/Users/rtaujale/Dropbox/Projects/collaborative/13C_mouse_INETA/raw_data/p
yINETA_AlaRef_2023/3.tilt.sym.ft2=CX1[Present](183.33,219.95):736758100000
.0;CX2[Present](36.39,220.0):1143852100000.0;"

Network27
"/Users/rtaujale/Dropbox/Projects/collaborative/13C_mouse_INETA/raw_data/p
yINETA_AlaRef_2023/3.tilt.sym.ft2=CX1[Present](36.72,221.34):2123371900000
.0;CX2[Present](184.84,221.45):806637000000.0;"

Network28
"/Users/rtaujale/Dropbox/Projects/collaborative/13C_mouse_INETA/raw_data/p
yINETA_AlaRef_2023/3.tilt.sym.ft2=CX1[Present](52.6,90.24):1226222500000.0
;CX2[Present](37.61,90.11):831301160000.0;"

Network29
"/Users/rtaujale/Dropbox/Projects/collaborative/13C_mouse_INETA/raw_data/p
yINETA_AlaRef_2023/3.tilt.sym.ft2=CX1[Present](37.96,214.07):9147714000000
.0;CX2[Present](176.08,214.05):1828480900000.0;CX4[Present](62.97,239.03):
1783091400000.0;"

Network30
"/Users/rtaujale/Dropbox/Projects/collaborative/13C_mouse_INETA/raw_data/p
yINETA_AlaRef_2023/3.tilt.sym.ft2=CX1[Present](49.98,88.1):11130542000000.
0;CX2[Present](38.15,88.13):9924519000000.0;"

Network31
```

```
"/Users/rtaujale/Dropbox/Projects/collaborative/13C_mouse_INETA/raw_data/p
yINETA_AlaRef_2023/3.tilt.sym.ft2=CX1[Present](41.77,97.25):1237302600000.
0;CX2[Present](55.36,97.25):804044200000.0;"

Network32
"/Users/rtaujale/Dropbox/Projects/collaborative/13C_mouse_INETA/raw_data/p
yINETA_AlaRef_2023/3.tilt.sym.ft2=CX1[Present](56.02,98.68):1177642100000.
0;CX2[Present](42.53,98.68):1676964700000.0;"

Network33
"/Users/rtaujale/Dropbox/Projects/collaborative/13C_mouse_INETA/raw_data/p
yINETA_AlaRef_2023/3.tilt.sym.ft2=CX1[Present](43.35,106.54):991751600000.
0;CX2[Present](63.12,106.42):1745454000000.0;"

Network34
"/Users/rtaujale/Dropbox/Projects/collaborative/13C_mouse_INETA/raw_data/p
yINETA_AlaRef_2023/3.tilt.sym.ft2=CX1[Present](53.69,97.62):763946140000.0
;CX2[Present](43.76,97.62):797534300000.0;"

Network35
"/Users/rtaujale/Dropbox/Projects/collaborative/13C_mouse_INETA/raw_data/p
yINETA_AlaRef_2023/3.tilt.sym.ft2=CX1[Present](44.0,218.17):2609389000000.
0;CX3[Present](174.37,218.7):938311750000.0;CX6[Present](59.26,233.4):8884
47960000.0;"

Network36
"/Users/rtaujale/Dropbox/Projects/collaborative/13C_mouse_INETA/raw_data/p
yINETA_AlaRef_2023/3.tilt.sym.ft2=CX1[Present](60.23,104.39):729235260000.
0;CX2[Present](44.13,104.39):2625134700000.0;"

Network37
"/Users/rtaujale/Dropbox/Projects/collaborative/13C_mouse_INETA/raw_data/p
yINETA_AlaRef_2023/3.tilt.sym.ft2=CX1[Present](180.62,225.68):736136270000
.0;CX2[Present](44.86,225.79):824767150000.0;"

Network38
"/Users/rtaujale/Dropbox/Projects/collaborative/13C_mouse_INETA/raw_data/p
yINETA_AlaRef_2023/3.tilt.sym.ft2=CX1[Present](178.15,223.78):181460910000
0.0;CX3[Present](46.0,223.94):4829206300000.0;"

Network39
"/Users/rtaujale/Dropbox/Projects/collaborative/13C_mouse_INETA/raw_data/p
yINETA_AlaRef_2023/3.tilt.sym.ft2=CX1[Present](46.86,115.12):1034964760000
.0;CX2[Present](49.38,96.25):714900200000.0;CX4[Present](68.23,115.23):722
591150000.0;"

Network40
"/Users/rtaujale/Dropbox/Projects/collaborative/13C_mouse_INETA/raw_data/p
yINETA_AlaRef_2023/3.tilt.sym.ft2=CX1[Present](52.76,114.37):1196998100000
.0;CX2[Present](61.89,114.52):1095730600000.0;"

Network41
```

```
"/Users/rtaujale/Dropbox/Projects/collaborative/13C_mouse_INETA/raw_data/p
yINETA_AlaRef_2023/3.tilt.sym.ft2=CX1[Present](53.12,230.86):1627207500000
.0;CX2[Present](177.71,230.86):915893500000.0;"

Network42
"/Users/rtaujale/Dropbox/Projects/collaborative/13C_mouse_INETA/raw_data/p
yINETA_AlaRef_2023/3.tilt.sym.ft2=CX1[Present](55.28,229.4):813073200000.0
;CX2[Present](173.87,229.56):1173121000000.0;"

Network43
"/Users/rtaujale/Dropbox/Projects/collaborative/13C_mouse_INETA/raw_data/p
yINETA_AlaRef_2023/3.tilt.sym.ft2=CX1[Present](177.5,233.48):787060900000.
0;CX2[Present](55.83,233.72):1235917300000.0;"

Network44
"/Users/rtaujale/Dropbox/Projects/collaborative/13C_mouse_INETA/raw_data/p
yINETA_AlaRef_2023/3.tilt.sym.ft2=CX1[Present](58.28,128.55):702064950000.
0;CX2[Present](70.17,128.58):787518400000.0;"

Network45
"/Users/rtaujale/Dropbox/Projects/collaborative/13C_mouse_INETA/raw_data/p
yINETA_AlaRef_2023/3.tilt.sym.ft2=CX1[Present](60.68,122.31):772474140000.
0;"

Network46
"/Users/rtaujale/Dropbox/Projects/collaborative/13C_mouse_INETA/raw_data/p
yINETA_AlaRef_2023/3.tilt.sym.ft2=CX1[Present](68.97,130.51):958458230000.
0;CX2[Present](62.07,130.8):1200856200000.0;CX3[Present](61.04,130.4):1019
011200000.0;"

Network47
"/Users/rtaujale/Dropbox/Projects/collaborative/13C_mouse_INETA/raw_data/p
yINETA_AlaRef_2023/3.tilt.sym.ft2=CX1[Present](175.75,238.53):780957060000
.0;CX2[Present](63.04,238.26):1752197800000.0;"

Network48
"/Users/rtaujale/Dropbox/Projects/collaborative/13C_mouse_INETA/raw_data/p
yINETA_AlaRef_2023/3.tilt.sym.ft2=CX1[Present](74.1,146.86):2497356800000.
0;CX2[Present](72.59,146.83):4374824700000.0;CX4[Present](63.49,137.64):36
40010000000.0;"

Network49
"/Users/rtaujale/Dropbox/Projects/collaborative/13C_mouse_INETA/raw_data/p
yINETA_AlaRef_2023/3.tilt.sym.ft2=CX1[Present](63.56,142.6):3657581700000.
0;CX2[Present](78.81,142.35):5037370000000.0;"

Network50
"/Users/rtaujale/Dropbox/Projects/collaborative/13C_mouse_INETA/raw_data/p
yINETA_AlaRef_2023/3.tilt.sym.ft2=CX1[Present](73.62,138.44):806795100000.
0;CX2[Present](64.94,138.43):1153838100000.0;"

Network51
```

```
"/Users/rtaujale/Dropbox/Projects/collaborative/13C_mouse_INETA/raw_data/p
yINETA_AlaRef_2023/3.tilt.sym.ft2=CX1[Present](65.74,243.31):759493750000.
0;CX2[Present](177.61,243.39):912987100000.0;"

Network52
"/Users/rtaujale/Dropbox/Projects/collaborative/13C_mouse_INETA/raw_data/p
yINETA_AlaRef_2023/3.tilt.sym.ft2=CX1[Present](85.82,153.46):718851300000.
0;CX2[Present](67.59,153.56):808582050000.0;"

Network53
"/Users/rtaujale/Dropbox/Projects/collaborative/13C_mouse_INETA/raw_data/p
yINETA_AlaRef_2023/3.tilt.sym.ft2=CX1[Present](73.8,142.03):982890840000.0
;CX2[Present](67.95,141.93):969492270000.0;"

Network54
"/Users/rtaujale/Dropbox/Projects/collaborative/13C_mouse_INETA/raw_data/p
yINETA_AlaRef_2023/3.tilt.sym.ft2=CX1[Present](171.19,239.93):722172400000
.0;CX2[Present](68.8,239.96):978746540000.0;"

Network55
"/Users/rtaujale/Dropbox/Projects/collaborative/13C_mouse_INETA/raw_data/p
yINETA_AlaRef_2023/3.tilt.sym.ft2=CX1[Present](71.04,255.69):2677792300000
.0;CX2[Present](184.53,255.64):1155736800000.0;"

Network56
"/Users/rtaujale/Dropbox/Projects/collaborative/13C_mouse_INETA/raw_data/p
yINETA_AlaRef_2023/3.tilt.sym.ft2=CX1[Present](94.87,169.46):2254432000000
.0;CX2[Present](74.43,150.18):1827269200000.0;CX3[Present](75.56,150.18):2
142712600000.0;"

Network57
"/Users/rtaujale/Dropbox/Projects/collaborative/13C_mouse_INETA/raw_data/p
yINETA_AlaRef_2023/3.tilt.sym.ft2=CX1[Present](78.72,155.7):5029263600000.
0;CX2[Present](98.79,175.98):3195300700000.0;CX3[Present](76.99,155.7):250
6849500000.0;"

Network58
"/Users/rtaujale/Dropbox/Projects/collaborative/13C_mouse_INETA/raw_data/p
yINETA_AlaRef_2023/3.tilt.sym.ft2=CX1[Present](77.39,167.15):853489000000.
0;CX2[Present](89.73,167.02):741308200000.0;"

Network59
"/Users/rtaujale/Dropbox/Projects/collaborative/13C_mouse_INETA/raw_data/p
yINETA_AlaRef_2023/3.tilt.sym.ft2=CX1[Present](79.69,251.91):721369000000.
0;CX2[Present](172.08,251.96):731503600000.0;"

Network60
"/Users/rtaujale/Dropbox/Projects/collaborative/13C_mouse_INETA/raw_data/p
yINETA_AlaRef_2023/3.tilt.sym.ft2=CX1[Present](85.51,177.14):669664540000.
0;CX2[Present](91.48,177.14):719250200000.0;"

Network61
```

```
"/Users/rtaujale/Dropbox/Projects/collaborative/13C_mouse_INETA/raw_data/p
yINETA_AlaRef_2023/3.tilt.sym.ft2=CX1[Present](168.57,273.8):726629500000.
0;CX2[Present](105.15,273.37):786833700000.0;"

Network62
"/Users/rtaujale/Dropbox/Projects/collaborative/13C_mouse_INETA/raw_data/p
yINETA_AlaRef_2023/3.tilt.sym.ft2=CX1[Present](126.48,244.29):787265360000
.0;CX2[Present](117.68,244.4):738509000000.0;"

Network63
"/Users/rtaujale/Dropbox/Projects/collaborative/13C_mouse_INETA/raw_data/p
yINETA_AlaRef_2023/3.tilt.sym.ft2=CX1[Present](119.39,254.13):741678700000
.0;CX2[Present](134.66,254.13):770011300000.0;"

Network64
"/Users/rtaujale/Dropbox/Projects/collaborative/13C_mouse_INETA/raw_data/p
yINETA_AlaRef_2023/3.tilt.sym.ft2=CX1[Present](121.12,272.98):734866500000
.0;CX2[Present](151.69,272.98):850254230000.0;"

Network65
"/Users/rtaujale/Dropbox/Projects/collaborative/13C_mouse_INETA/raw_data/p
yINETA_AlaRef_2023/3.tilt.sym.ft2=CX1[Present](126.82,281.2):1024876700000
.0;CX2[Present](154.51,281.4):980925200000.0;"

Network66
"/Users/rtaujale/Dropbox/Projects/collaborative/13C_mouse_INETA/raw_data/p
yINETA_AlaRef_2023/3.tilt.sym.ft2=CX1[Present](151.08,278.03):750467500000
.0;CX2[Present](126.96,278.13):1001294860000.0;"

Network67
"/Users/rtaujale/Dropbox/Projects/collaborative/13C_mouse_INETA/raw_data/p
yINETA_AlaRef_2023/3.tilt.sym.ft2=CX1[Present](178.26,363.89):185129360000
0.0;CX2[Present](186.07,363.89):793620840000.0;"
```

Supplementary Information 6: Evaluation of the consistency of annotation between the original INETA and PyINETA

We evaluated the performance of our new Python pipeline. For this evaluation, we used the same INADEQUATE spectra that were used in the original study of INETA[1] as input files for PyINETA. They are INADEQAUATE spectra collected for endo- and exometabolites of $^{13}C$-labeled *Caenorhabditis elegans*. The results were consistent (Supplementary Table 7); out of 29 metabolites annotated in the previous study, 26 were found in the new pipeline. They covered a range of compounds, including amino acids, an amino acid derivative (creatine), organic acids, a fatty acid (palmitic acid), a polyamine (putrescine), a pyrimidine derivative (uracil), and sugar derivatives. Three of them that were not included in our results are *N*-acetylglycine, stearic acid, and glutathione oxidized.

Supplementary Table S1. List of adjustable parameters in PyINETA. Those parameters are contained in each configuration file.

| Parameter | Function | Value used for the mouse study | Note |
|---|---|---|---|
| ***For INADEQUATE*** | | | |
| *Peak picking* | | | |
| PPmin | Peak intensity minimum threshold | 5.7e10 | Clendinen et al. (2015) |
| PPmax | Peak intensity minimum threshold | 4e11 | Clendinen et al. (2015) |
| steps | Number of iterations | 10 | This study |
| PPCS | Chemical shift threshold for clustering | 1 | Clendinen et al. (2015) |
| PPDQ | Double quantum threshold for clustering | 2 | Clendinen et al. (2015) |
| *Network finding* | | | |
| DQT | Double quantum tolerance | 0.2 ppm | Clendinen et al. (2015) |
| SumXY | Sum tolerance | 2 | This study |
| SDT | Symmetrical/diagonal tolerance | 0.5 ppm | Clendinen et al. (2015) |
| CST | Chemical shift tolerance for vertical connection | 0.05 ppm | Clendinen et al. (2015) |
| *Database matching* | | | |
| Ambiguity | Ambiguity allowance | 1 | Clendinen et al. (2015) |
| CSMT | Chemical shift match tolerance | 1 ppm | Clendinen et al. (2015) |
| NCMT | Peak match tolerance | 2 | Clendinen et al. (2015) |
| DQMT | Double quantum threshold | 4 | Clendinen et al. (2015) |
| ***For JRES*** | | | |
| Peak_Width_1D | JRES peak width to be used to calculate peak area | 0.5 ppm | This study |
| Intensity_threshold_1D | Intensity threshold to define presence or absence of peak | 10,000 | This study |

Supplementary Table S2. Composition of the diet that was fed to the three mice in this study. For details on the composition of bacterial $^{13}C$ protein hydrolysate, see Supplementary Table S3.

| **Component** | **g/kg** |
|---|---|
| Sucrose | 350.0 |
| Bacterial $^{13}C$ protein hydrolysate | 200.0 |
| Maltodextrin | 130.0 |
| Corn Starch | 100.0 |
| Soybean Oil | 80.0 |
| Cellulose | 72.2 |
| Mineral Mix, AIN-93M-MX (94049) | 45.0 |
| Vitamin Mix, AIN-93-VX (94047) | 14.0 |
| L-Cystine | 3.0 |
| Calcium Phosphate, dibasic | 3.0 |
| Choline Bitartrate | 2.5 |
| Ferric Citrate | 0.25 |
| THBQ, antioxidant | 0.024 |

Supplementary Table S3. Amino acid composition of the hydrolysate that was fed to the three mice in this study (See also for Supplementary Table S2). The data are from a product sheet from Silantes. *: essential amino acid

| **Amino acid** | **Percent** |
|---|---|
| Asp | 20.56 |
| Thr* | 3.70 |
| Ser | 4.10 |
| Glu | 11.10 |
| Gly | 11.10 |
| Ala | 14.8 |
| Val* | 2.98 |
| Met* | 1.78 |
| Ileu* | 2.10 |
| Leu* | 5.26 |
| Tyr | 1.80 |
| Phe* | 2.40 |
| His* | 9.65 |
| Lys* | 3.38 |
| Arg | 2.30 |
| Pro | 2.99 |

Supplementary Table S4. Amount of the samples used for the mouse samples in this study. The weight is based on dry weight.

| Tissue | Mouse ID | Weight or volume |
|---|---|---|
| Liver | 1 | 1092 mg |
| | 2 | 998.9 mg |
| | 3 | 1000 mg |
| Adrenal Gland | 1 | 32.5 mg |
| | 2 | 22.3 mg |
| | 3 | 36.6 mg |
| Lung – Only 75% of lung tissue | 1 | 203.1 mg |
| | 2 | 353.9 mg |
| | 3 | 171.2 mg |
| Muscle – Only 50% of muscle tissue | 1 | 121.1 mg |
| | 2 | 137.6 mg |
| | 3 | 82.7 mg |
| Pancreas | 1 | 162.4 mg |
| | 2 | 152.6 mg |
| | 3 | 165.9 mg |
| Plasma | 1 | 50 µL |
| | 2 | 50 µL |
| | 3 | 50 µL |
| Brain | 1 | 226.5 mg |
| | 2 | 204.3 mg |
| | 3 | 226.5 mg |
| Spleen | 1 | 155.7 mg |
| | 2 | 154.8 mg |
| | 3 | 184.3 mg |
| Thymus | 1 | 59.83 mg |
| | 2 | 105.6 mg |
| | 3 | 68.3 mg |

Supplementary Table S5. Parameters used for NMR experiments. SW, spectral width; TD, size of FID; NS, number of scans.

| Figure | Experiment | Pulse program | SW (ppm) | | TD | | NS | Note |
|---|---|---|---|---|---|---|---|---|
| | | | F2 | F1 | F2 | F1 | | |
| 2 | INADEQUATE | INADEQUATEAD | 202 | 404 | 4096 | 2048 | 4 | |
| 3 | INADEAUATE | inadphppsp | 214 | 433 | 4096 | 2048 | 16 | |
| 3 | JRES | jresdcqf | 214 | 0.66 | 32768 | 50 | 64 | The original uniform 180$^{o}$ pulse was replaced with an adiabatic pulse |

Supplementary Table S6. Summary of NMRPipe processing parameters used for INADEQUATE and JRES experiments. SP: adjustable sine window function, ZF: zero filling. Original NMRPipe scripts are also available in Metabolomics Workbench.

| Figure | Experiment | SP<br>F2 and F1 | ZF<br>F2 and F1 | Baseline correction<br>F2 and F1 | Note |
|---|---|---|---|---|---|
| 2 | INADEQUATE | -off 0.0 -end 0.98 -pow 2 | -auto | POLY -auto | Linear prediction (LP -fb) was additionally used for F1 |
| 3 | INADEQUATE | -off 0.0 -end 0.95 -pow 2 | -auto | MED | – |
| 3 | JRES | -off 0.0 -end 0.98 -pow 2 | -auto | – | Magnitude calculation (MC) was additionally used; spectra were further tilted and symmetrized |

Supplementary Table S7. List of metabolites found by PyINETA for endo- and exometabolites from *Caenorhabditis elegans*. See also Experimental Section for the details on the confidence scores.

| Compound name | Ambiguity score | Hit score | Coverage Score | Type |
|---|---|---|---|---|
| ***Endometabolites*** | | | | |
| Proline | 0 | 0.25 | 1 | Found in both the previous and this studies |
| Valeric acid | 0 | 0.25 | 1 | Found in both the previous and this studies |
| Glucose 1,6 bisphosphate | 0 | 0.2 | 1 | Found in both the previous and this studies |
| Histidine | 0.333 | 0.25 | 1 | Found in both the previous and this studies |
| Threonine | 0 | 0.333 | 1 | Found in both the previous and this studies |
| Arabitol | 0 | 0.25 | 1 | Found in both the previous and this studies |
| Alanine | 0 | 0.5 | 1 | Found in both the previous and this studies |
| Isoleucine | 0 | 0.2 | 1 | Found in both the previous and this studies |
| Propionic acid | 0 | 0.5 | 1 | Found in both the previous and this studies |
| 3-hydroxybutyrate | 0 | 0.333 | 1 | Found in both the previous and this studies |
| Lactic acid | 0 | 0.5 | 1 | Found in both the previous and this studies |
| Valine | 0.4 | 0.25 | 1 | Found in both the previous and this studies |
| Palmitic acid | 0 | 0.2 | 0.625 | Found in both the previous and this studies |
| ***Exometabolites*** | | | | |
| Isoleucine | 0 | 0.2 | 1 | Found in both the previous and this studies |
| Alanine | 0 | 0.5 | 1 | Found in both the previous and this studies |
| Threonine | 0 | 0.333 | 1 | Found in both the previous and this studies |
| Glycine | 0 | 1 | 1 | Found in both the previous and this studies |
| Proline | 0 | 0.25 | 1 | Found in both the previous and this studies |
| Ornithine | 0 | 0.25 | 1 | Found in both the previous and this studies |
| Serine | 0 | 0.5 | 1 | Found in both the previous and this studies |
| Glucose 1,6 bisphosphate | 0 | 0.2 | 1 | Found in both the previous and this studies |
| Lysine | 0 | 0.2 | 1 | Found in both the previous and this studies |

| | | | | |
|---|---|---|---|---|
| Glutamine | 0 | 0.25 | 1 | Found in both the previous and this studies |
| Lactic acid | 0 | 0.5 | 1 | Found in both the previous and this studies |
| Glucoronate | 0 | 0.5 | 0.75 | Found in both the previous and this studies |
| Putrescine | 0 | 0.333 | 1 | Found in both the previous and this studies |
| Acetic acid | 0 | 1 | 1 | Found in both the previous and this studies |
| Arabitol | 0 | 0.25 | 1 | Found in both the previous and this studies |
| Creatine | 0 | 1 | 1 | Found in both the previous and this studies |
| Valine | 0.4 | 0.25 | 1 | Found in both the previous and this studies |
| Allantoin | 0 | 1 | 0.667 | Found in both the previous and this studies |
| Propionic acid | 0 | 0.5 | 1 | Found in both the previous and this studies |
| Succinic acid | 0 | 0.333 | 1 | Found in both the previous and this studies |
| Methionine | 0 | 0.333 | 0.5 | Found in both the previous and this studies |
| Uracil | 0 | 0.5 | 1 | Found in both the previous and this studies |

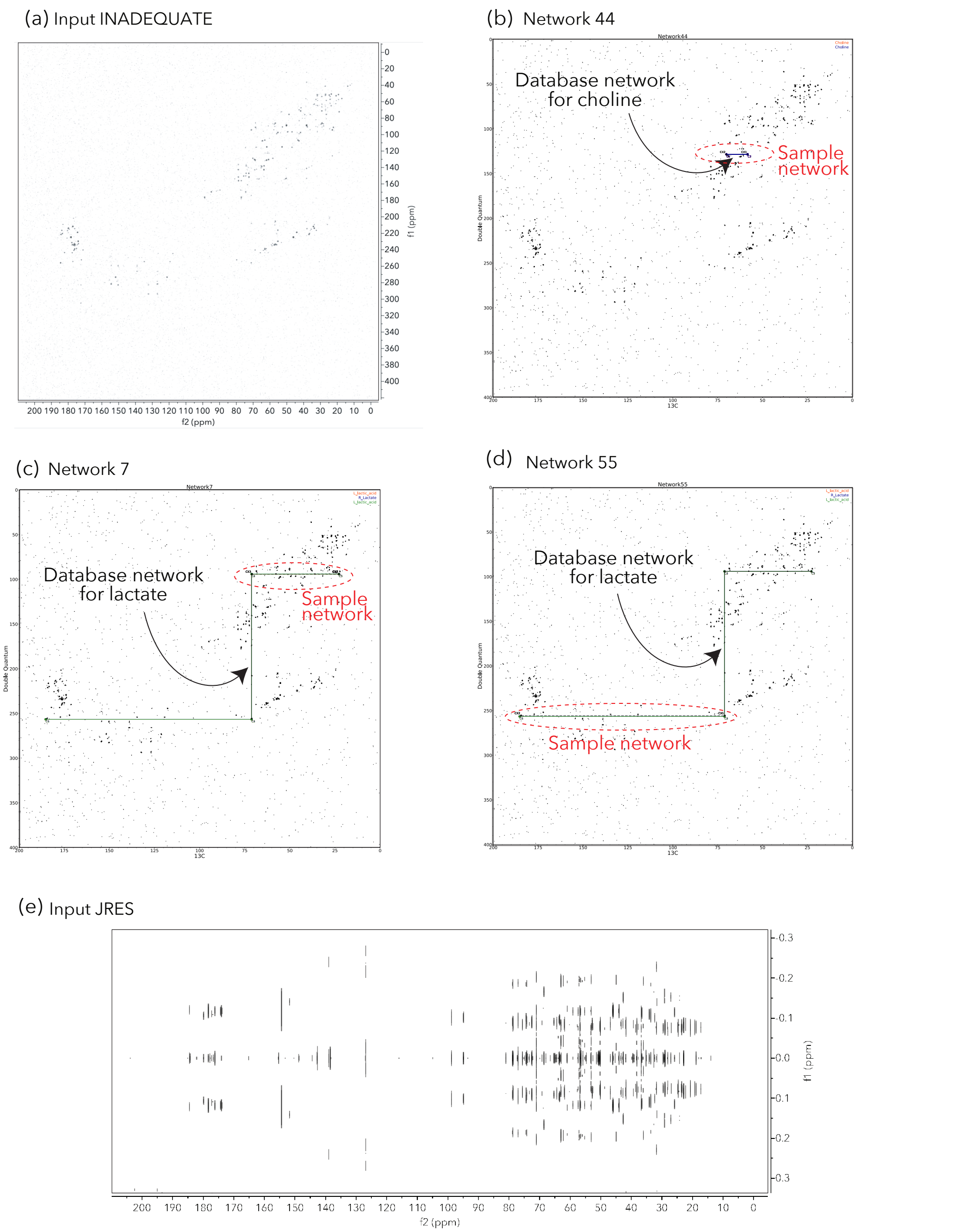


Supplementary Figure S1 (a) Input INADEQUATE spectrum for the liver sample; (b) Network 44 of PyINETA; (c) Network 7; (d) Network 55; (e) Input JRES spectrum for the liver sample. For a and e, the plots were produced by MestReNova v14.3.0. For b, c, and d, the plots are the original outputs from PyINETA, with a slight graphical modification.

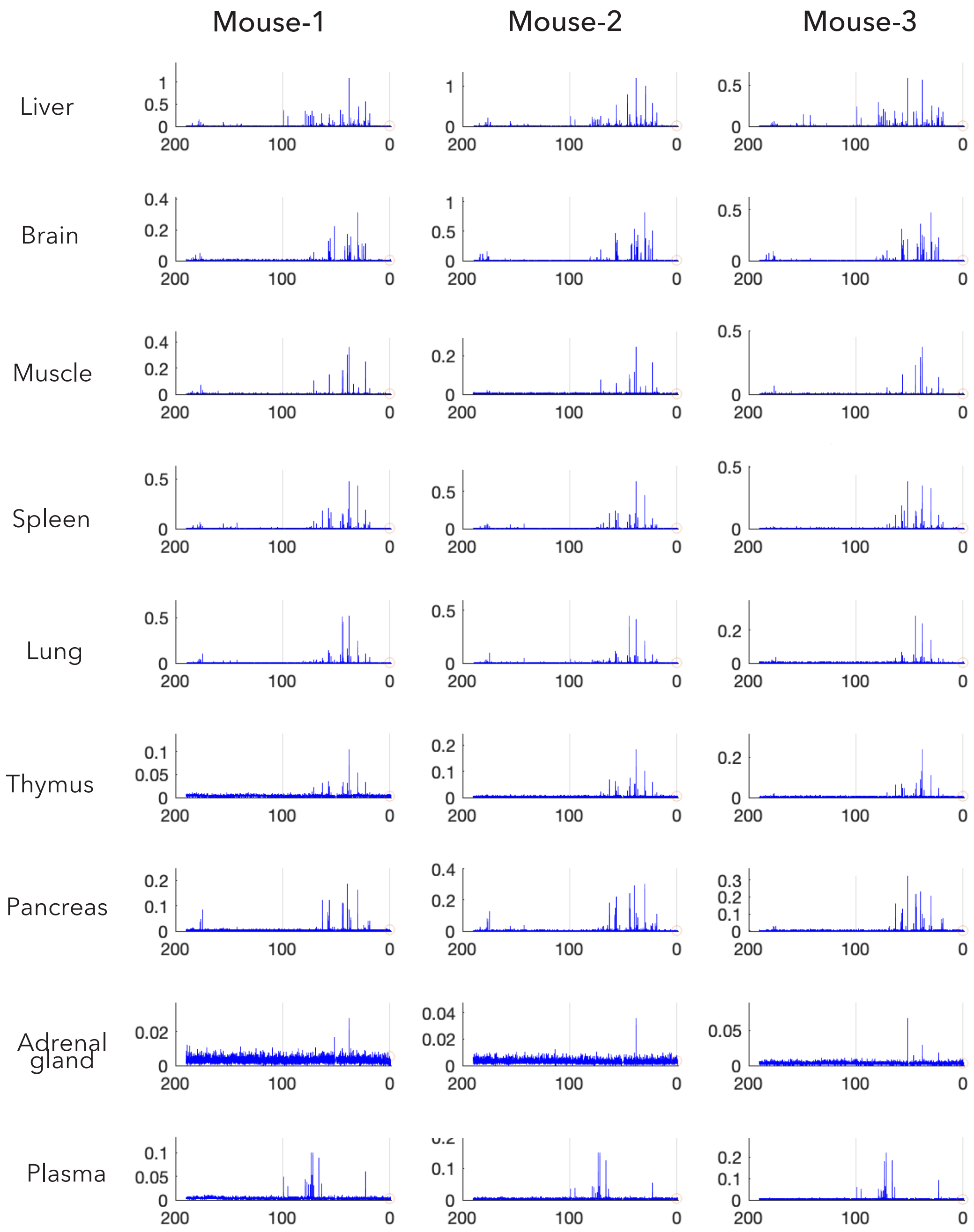


Supplementary Figure S2 JRES projection spectra for the mouse tissues. Solvent MeOH peaks around 50 ppm are removed and not shown.

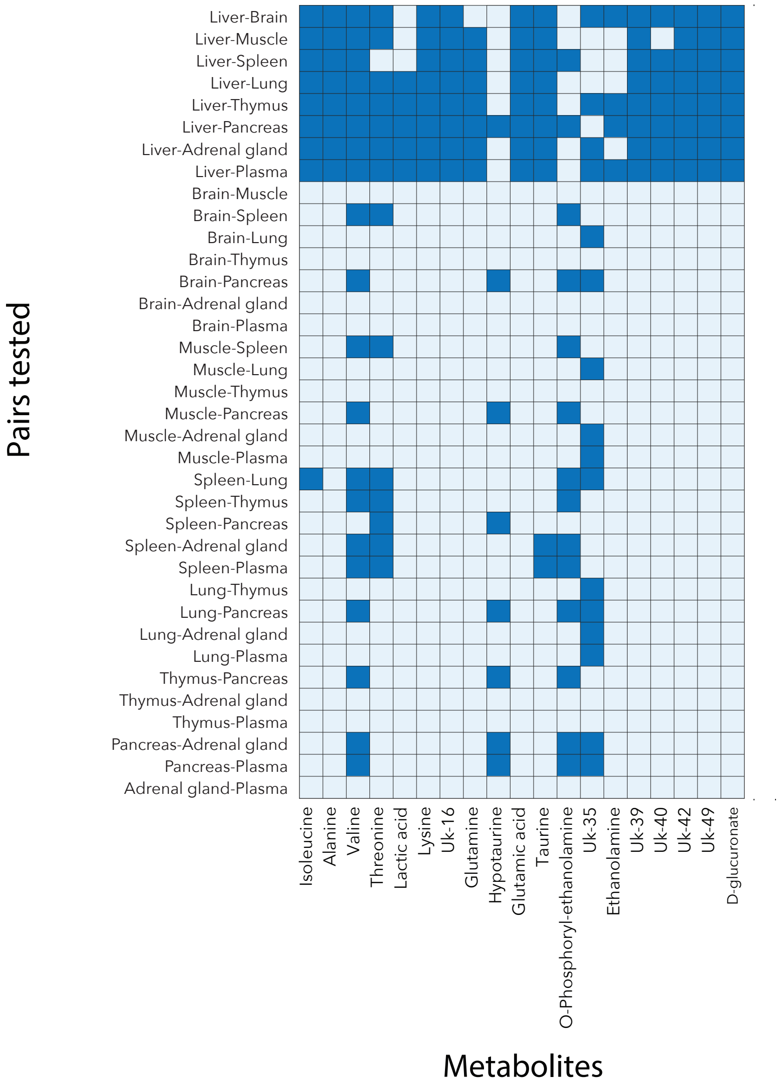


Supplementary Figure S3 Results of ANOVA and multiple comparison. Metabolites intensities were compared between tissues for the three mice. When there is a significant difference between tissues, they are highlighted in dark blue (*p* < 0.05 with Bonfferoni Correction). This information is summarized in Figure 3.

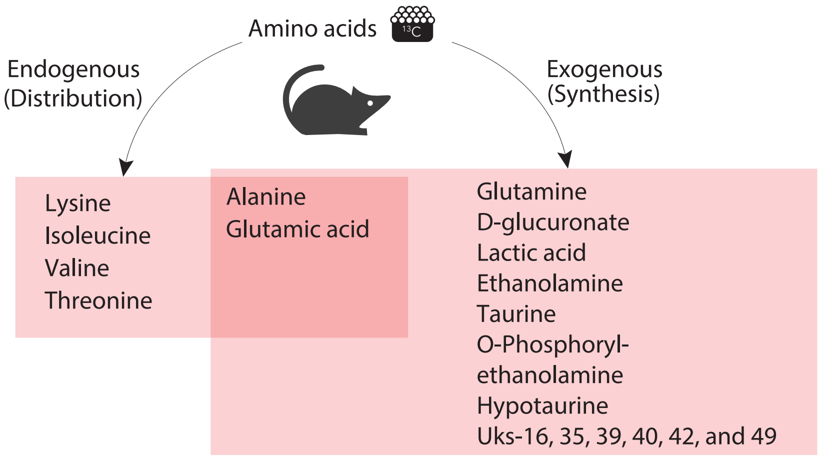


Supplementary Figure 4 Possible distribution mechanisms of metabolites in this study

**Reference**

1. Clendinen, C. S.; Pasquel, C.; Ajredini, R.; Edison, A. S., 13C NMR Metabolomics: INADEQUATE Network Analysis. *Anal Chem* **2015,** *87* (11), 5698-706.
